# Widespread transfer of mobile antibiotic resistance genes within individual gut microbiomes revealed through bacterial Hi-C

**DOI:** 10.1101/2020.03.19.998526

**Authors:** Alyssa Kent, Albert Vill, Qiaojuan Shi, Michael J. Satlin, Ilana Lauren Brito

## Abstract

The gut microbiome harbors a ‘silent reservoir’ of antibiotic resistance (AR) genes that is thought to contribute to the emergence of multidrug-resistant pathogens through the process of horizontal gene transfer (HGT). To counteract the spread of AR genes, it is paramount to know which organisms harbor mobile AR genes and with which organisms they engage in HGT. Despite methods to characterize the bulk presence^1^, abundance^2^ and function^3^ of AR genes in the gut, technological limitations of short-read sequencing have precluded linking bacterial taxa to specific mobile genetic elements (MGEs) and their concomitant AR genes. Here, we apply and evaluate a high-throughput, culture-independent method for surveilling the bacterial carriage of MGEs, based on bacterial Hi-C protocols. We compare two healthy individuals with a cohort of seven neutropenic patients undergoing hematopoietic stem cell transplantation, who receive multiple courses of antibiotics throughout their prolonged hospitalizations, and are thus acutely vulnerable to the threat of multidrug-resistant infections^4^. We find that the networks of HGT are surprisingly distinct between individuals, yet AR and mobile genes are more dispersed across taxa within the neutropenic patients than the healthy subjects. Our data further suggest that HGT is occurring throughout the course of treatment in the microbiomes of neutropenic patients and within the guts of healthy individuals over a similar timeframe. Whereas most efforts to understand the spread of AR genes have focused on pathogenic species, our findings shed light on the role of the human gut microbiome in this process.

---

The acquisition of antibiotic resistance (AR) genes has rendered important pathogens, such as multidrug-resistant (MDR) Enterobacteriaceae and *Pseudomonas aeruginosa*, nearly or fully unresponsive to antibiotics. It is widely accepted that these so-called ‘superbugs’ acquire AR genes through the process of horizontal gene transfer (HGT) with members of the human microbiome with whom they come into contact^5^. The emergence of these MDR bacteria threatens our ability to perform life-saving interventions, such as curative hematopoietic cell transplants for patients with hematologic malignancies^6^. Furthermore, antibiotic use, required for vital prophylaxis in these patients, has been proposed as a trigger for HGT. Although tools are available to identify AR genes within the gut microbiome, and characterize their function^7^, abundance^8,9^ and their host-associations^10^, no studies have attempted to monitor the bacterial host associations of AR genes and mobile elements during relatively short periods, such as during these patients’ hospitalizations.

To determine the bacterial hosts of mobile AR genes, we utilized a high-throughput chromatin conformation capture (Hi-C) method aimed at sampling long-range interactions within single bacterial genomes^11,12,13^. Briefly, while cells are still intact, DNA within individual cells is crosslinked by formaldehyde. Cells are then lysed and the DNA is cut with restriction enzymes, biotinylated, and subjected to dilute ligation to promote intra-molecular linkages between crosslinked DNA. Crosslinking is reversed and then ligated DNA molecules are pulled-down and made into DNA libraries for sequencing. As is, this protocol has been used to improve metagenomic assemblies of bacterial genomes^14^ and has identified a handful of strong plasmid- and phage-bacterial host associations^15,16,17,18^, suggesting that this technique could be applied to link mobile genes with specific taxa more broadly and to observe the process of HGT over time.

Here, we develop a modified version of current Hi-C protocols and analytical pipelines (Extended Data Figure 1) in conjunction with metagenomic shotgun sequencing to surveil the bacterial taxa harboring specific mobile AR genes in the gut microbiomes of two healthy individuals and seven patients undergoing hematopoietic stem cell transplantation. These patients have prolonged hospitalizations during their transplant (21 ± 4 days) and often receive multiple courses of antibiotic therapy, increasing the likelihood of an MDR infection. As a result of their condition and treatment, these patients face mortality rates of 40-70% when bacteremic with carbapenem-resistant Enterobacteriaceae (CRE) or carbapenem-resistant *Pseudomonas aeruginosa*^19^, and therefore represent a salient population for surveillance and one in which MDR pathogens may emerge and/or amplify under antibiotic selection. Gut microbiome samples for patients and healthy subjects were collected over a 2-3-week period, which, for the neutropenic patients started upon admission for transplant and continued during their hospitalization until neutrophil engraftment (Figure 1A, Extended Data Table 1,2).

**Figure 1.**
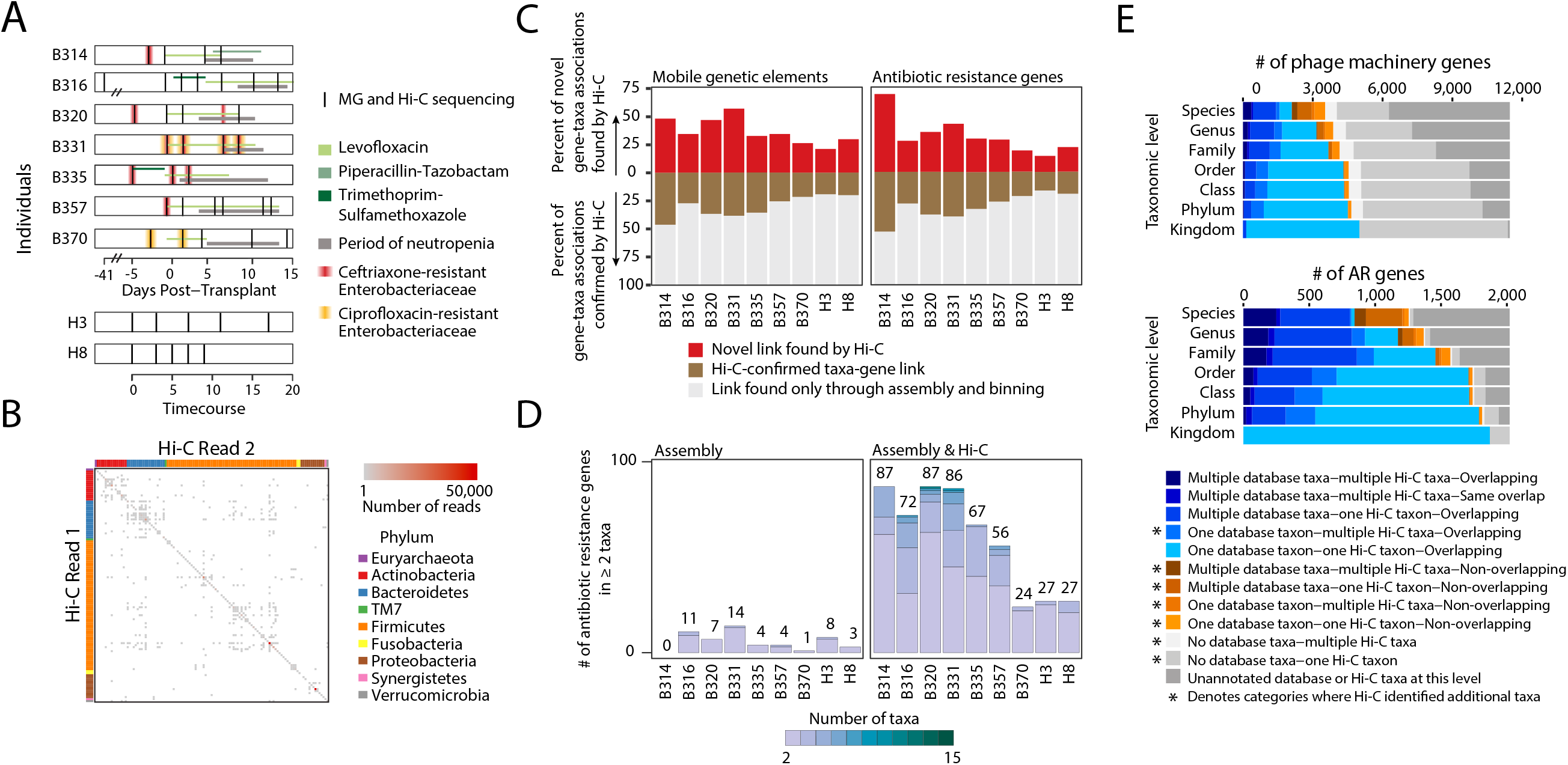
Hi-C can be used to track mobile genetic elements. (A) Neutropenic hematopoetic stem cell transplant (B) recipients’ and healthy (H) individuals’ timecourses included in the study are depicted, with periods of neutropenia (gray) and antibiotic use (green). Black lines indicate time points for which metagenomic and Hi-C libraries were constructed. Red lines indicate gastrointestinal colonization with MDR enteric pathogens. (B) Each Hi-C read pair that maps to two non-mobile contigs is plotted according to the taxonomic assignment of each read. Color depicts the number of reads linking contigs according to taxonomy. (C) The percent of the total taxa-mobile gene (left) and taxa-AR gene (right) associations observed from metagenomic assembly that are supported by two or more Hi-C links (brown) is plotted, along with the percent additional interactions gained by using Hi-C (red). (D) Stacked barplots showing the number of species-level taxa to which each AR gene (clustered at 99% identity) is assigned within each patient, and across patients. Only those genes assigned to 2 or more taxa are shown. We either used metagenomic assemblies alone to assign taxonomies (left) or combined with Hi-C libraries considering those taxa-gene assignments with evidence from at least two Hi-C reads. The numbers above each stacked barplot represent the total number of AR genes with 2 or more taxonomic associations. (E) Horizontal stacked bar plots show the percentage of unique phage genes (defined as 95% similar) (above) or AR genes (below) in the metagenomic assemblies and the origins of their taxonomic associations, identified either by BLAST to NCBI’s NT (for phage) or PATRIC’s reference database (for AR genes) or through Hi-C linkages.

We introduced a number of improvements to current bacterial Hi-C protocols. We changed sample storage and optimized restriction enzymes to improve the congruence between the composition of metagenomic and Hi-C sequencing libraries (Extended Data Figure 2). We also integrated Nextera XT sequencing library preparation directly into the Hi-C experimental protocol, streamlining operations and decreasing sample preparation time. Importantly, within diverse bacterial communities such as the gut microbiome, MGEs may be highly promiscuous and recombinogenic, complicating both assembly^20^ and linkage analyses^21^. Therefore, we implemented a computational workflow to assemble genomes, separating large integrated MGEs onto their own contigs, and allowing them to associate with genomes via binning or Hi-C connections. In a mock community of three organisms, each harboring an identifiable plasmid, we are able to confidently link each plasmid to its nascent genome (Extended Data Figure 3).

Our Hi-C experimental and computational approach results in robust linkages between contigs in human microbiome samples. Hi-C read pairs linking non-mobile contigs with contradictory taxonomic annotations are rarely observed (3.4% at the genus-level) and likely represent homologous sequence matches, highlighting the purity of our Hi-C libraries (Figure 1B). Hi-C read pairs linking two contigs are preferentially recruited to contigs that are longer and more abundant, but to a lesser degree than expected, reducing potential bias in our dataset toward highly abundant organisms (Extended Data Figure 4). We binned contigs using several tools (Maxbin^22^, MetaBat and Concoct), and applied a binning aggregation strategy, DAS Tool^23^, to obtain a set of draft genomic assemblies. As misassembly can resemble HGT, we removed assemblies with greater than 10% contamination, as determined by CheckM, resulting in taxonomically coherent assemblies (Extended Data Figure 5), albeit a greater number of unbinned contigs (24.6% of the total) (Extended Data Table 3). We then apply conservative criteria to link mobile and mobile AR-containing contigs with the genomic draft assemblies, considering an MGE part of a genome assembly only if it is directly linked to it by at least two uniquely-mapped Hi-C read-pairs. As MGEs are known to recombine, this mitigates the potential for falsely linking contigs that merely share common mobile genes. However, this also potentially reduces our ability for overall detection, especially for larger MGEs, since mobile contigs are often fragmented in metagenomic assemblies^24^. Nevertheless, we restricted our analysis to those AR-organism and MGE-organism linkages derived from high-confidence read mappings.

Hi-C significantly improved our ability to detect mobile gene-bacterial host linkages beyond standard metagenomic assembly alone. Hi-C confirms many of the AR gene-taxa and mobile gene-taxa associations observed in the metagenomic assemblies (30.49% ± 11.49% of the AR genes; 30.1% ± 9.52% of the mobile genes), but importantly adds on average 31.81% ± 16.28% AR gene associations and 36.64% ± 11.56% mobile gene associations to those observed by metagenome assembly alone (Figure 1C). Furthermore, whereas metagenomic assembly methods can generally link a single mobile gene cluster to one or two organisms, our Hi-C method was able to identify up to 15 bacterial hosts harboring the same AR or mobile gene, requiring two or more Hi-C linkages within a single individual (mean = 3.53 ± 5.69 bacterial hosts per AR gene, Figure 1D; mean = 6.85 ± 10.88 bacterial hosts per gene, Extended Data Figure 6). A larger percentage of AR and mobile genes overall (8.1%±5.2% vs. 0.9%±0.9% for AR genes and 6.1%±4.8% vs. 1.9%±1.7% for mobile genes) can be assigned to multiple taxonomies. These results were consistent with more stringent thresholds for Hi-C associations (Extended Data Figure 7).

Our data increases mobile and AR gene-taxa assignments above those observed using publicly available reference genomes, while focusing on those immediately relevant to the individual patient. We first investigated phage-host associations identified through Hi-C and compared them with those in NCBI, as many phage are host-specific^25^. Indeed, 43.5% of the phage genes with Hi-C genera-level assignments recapitulate known interactions (Figure 1E). However, broader genera-level associations are obtained for 64.2% of the unique phage genes in our database, reflecting apparent selection biases within our reference databases and the promiscuity of certain phage^26,27,28^. A greater percentage of AR genes with Hi-C genera-level taxonomic assignments, 82.8%, were evident in reference genomes. Yet, Hi-C expands genera-level assignments for 37.6% of the AR genes. Despite having a limited number of reads linking each mobile or AR gene to a particular taxa, our annotations are supported by the fact that Hi-C reads preferentially map near to these genes on the overall contig (Extended Data Figure 8).

We next sought to determine the extent to which we could capture associations using Hi-C. First, we performed a modified rarefaction analysis to determine whether the number of AR gene-taxa associations and mobile gene-taxa associations saturated with increased sequencing depth of our Hi-C libraries. Most of our samples saturated within our target sequencing depth (roughly 15 million paired reads), and sequencing samples to roughly four-fold this amount did not significantly increase the number of gene-taxa associations (Extended Data Figure 9). The number of contigs that recruited Hi-C reads (on average 18.3 ± 10.9%) was not dependent on sequencing depth, yet 88.2% ± 9.5 of our genome bins recruited two or more Hi-C reads, which amounts to 90.7% ± 9.0% of the taxa recruiting reads. This breadth is supported by the congruence of Hi-C libraries and metagenomic libraries (Extended Data Figure 2). We suspect that the variation in recruitment of Hi-C reads across the genome reflects either recurrent structural patterning of DNA^29^, differences in DNA-binding proteins available for cross-linking, and the distribution of restriction enzyme cut sites^30^. We next measured our ability to detect the same mobile genes across timepoints. If we consider only the AR gene-taxa associations we observe at least once, and we conservatively assume that should continue to observe the gene-taxa association, *i.e.* that the lack of repeated observation was due to the stochastic sampling process of Hi-C rather than HGT, we repeatedly detect an average of 66% of all possible associations where both the organism and AR genes were detectable in the draft assemblies but were not linked through Hi-C (Extended Data Figure 10).

Overall, within each person’s microbiome, mobile genes, including AR genes and HGT machinery genes, were distributed across a wide range of taxa (Extended Data Figures 11 and 12). Less than 10% of unique mobile genes and 19% of unique AR genes (clustered at 99% identity) were found across multiple patients, a finding consistent with previous surveys of MGEs across individuals^31^, indicating limited inter-personal or nosocomial transmission. Furthermore, for these MGEs found across patients, few of their host associations were conserved. We speculate that HGT may result in their dispersal within individual’s gut microbiomes and that selection may affect MGE-taxa associations at the level of individuals^32^. Despite heavy administration of antibiotics, the abundances of AR genes, even those conferring resistance to administered antibiotics, did not correspond with patient-specific therapeutic courses, a finding consistent with other patient-timecourses of mobile AR genes^33^, and possibly reflective of the low plasmid-based resistance to levofloxacin or combination antibiotic therapies (Extended Data Figure 13).

When comparing the networks of HGT within each individual’s gut microbiome, we expected to observe a strong preference for gene exchange between more closely related organisms, as previously observed when comparing exchange networks using reference genomes^34^. Whether HGT occurs more frequently in an individual’s gut is an essential question to understand the development and maintenance of the reservoir of AR genes in the gut microbiome, yet it has been difficult to answer for technical reasons. Using Hi-C, we find that the spread of AR genes and other mobile genes is significantly higher within an individual’s gut microbiome than between different individuals’ gut microbiomes (Figure 2A, Extended Data Figure 14). Beyond closely related pairs of organisms, there was considerable variation in the networks of shared AR and mobile genes across individuals (Extended Data Figures 15). Despite this, we find that those microbiomes similar in composition shared more of the same connections among the organisms present in both microbiomes (Figure 2B), most notably between the two healthy individuals.

**Figure 2.**
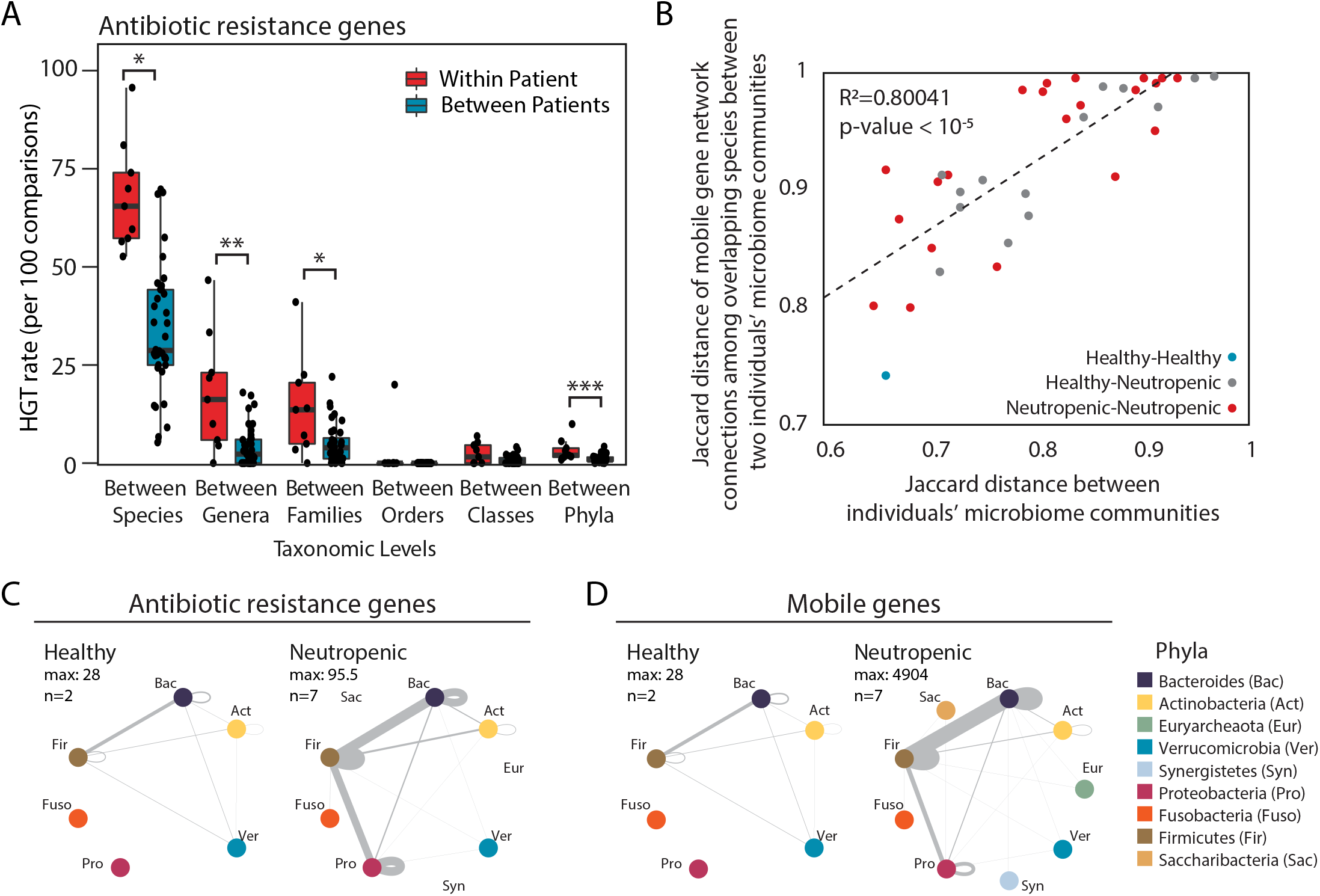
Networks of bacterial HGT are unique to each individuals’ gut microbiomes. (A) HGT rates (per 100 comparisons) of AR genes between organisms within each individual versus between individuals, according to those that share the same species, genus, family, order, class, and phylum are plotted for comparison. Significance was measured with Mann Whitney U-tests (two-sided; *, p<0.05; **, p<0.01; ***, p<0.005, ****, p<0.001;*****, p<0.0005). Boxplot represents the interquartile range where ends of the whiskers represent ±1.5*interquartile range and median value is indicated. (B) For each pair of individuals, the Jaccard distance of their composite microbiome compositions are plotted against the average Jaccard distance of the HGT network connections of mobile genes exchanged between organisms present in both individuals. Points are colored according to the health status of the donors being compared. (C) Network plots showing bacterial AR gene exchange according to phyla within healthy (left) and neutropenic (right) individuals’ microbiomes. n refers to the number of people included in the plot. (D) Network plots showing bacterial mobile gene exchange according to phyla within healthy (left) and neutropenic (right) individuals’ microbiomes. n refers to the number of people included in the plot.

Given their clinical importance, we focused on the gene-sharing networks of Proteobacteria, and more specifically, Enterobacteriaceae. Within all patients, gene exchange was most frequent within members of the same phylum (Figure 2C,D). In neutropenic patients, Proteobacteria shared genes outside their phylum most often with Firmicutes. The main transfer partners with Enterobacteriaceae were different across patients, but notably included both opportunistic pathogens (*i.e. Veillonella parvula* and *Enterococcus faecium*), commensals that may flourish post-antibiotic use (*i.e. Erysipelotrichaceae* sp.^35^), and even those organisms that have been considered as probiotic (*i.e. Faecalibacterium prausnitzii*^36^ and *Roseburia intestinalis*^37^).

Antibiotic treatment^38^ and inflammation^39^ are putative triggers for HGT, through the production of reactive oxygen species and DNA damage. We hypothesized that mucositis caused by cytotoxic chemotherapy, along with the selective pressures imposed by antibiotics and inflammation, would create conditions amenable to HGT in these neutropenic patients. We noticed that the average density of connections (percentage of actual connections of the total possible connections) between taxa and AR or mobile genes is greater in the neutropenic patients than the healthy individuals (Figure 3A). Several patients, B316, B320, B335 and B370, experienced increases in the proportion of overall gene-taxa connections, referred to as network density, during their timecourses. This was unrelated to the abundance of Enterobacteriaceae in the samples, which have been proposed as mediators of HGT^40^, the total abundance of AR genes, or the number of Hi-C reads (Extended Data Figure 16). Rather, we found that the only correlate was the number of taxa in a sample: as patients’ microbiomes became less diverse, the gene-taxa network density increased (Figure 3B). We hypothesize that this is caused either by an undefined selective pressure acting to preserve more connected organisms; or that once selection has occurred, organisms in less diverse populations will have increased contact rates and therefore greater opportunity for transfer.

**Figure 3.**
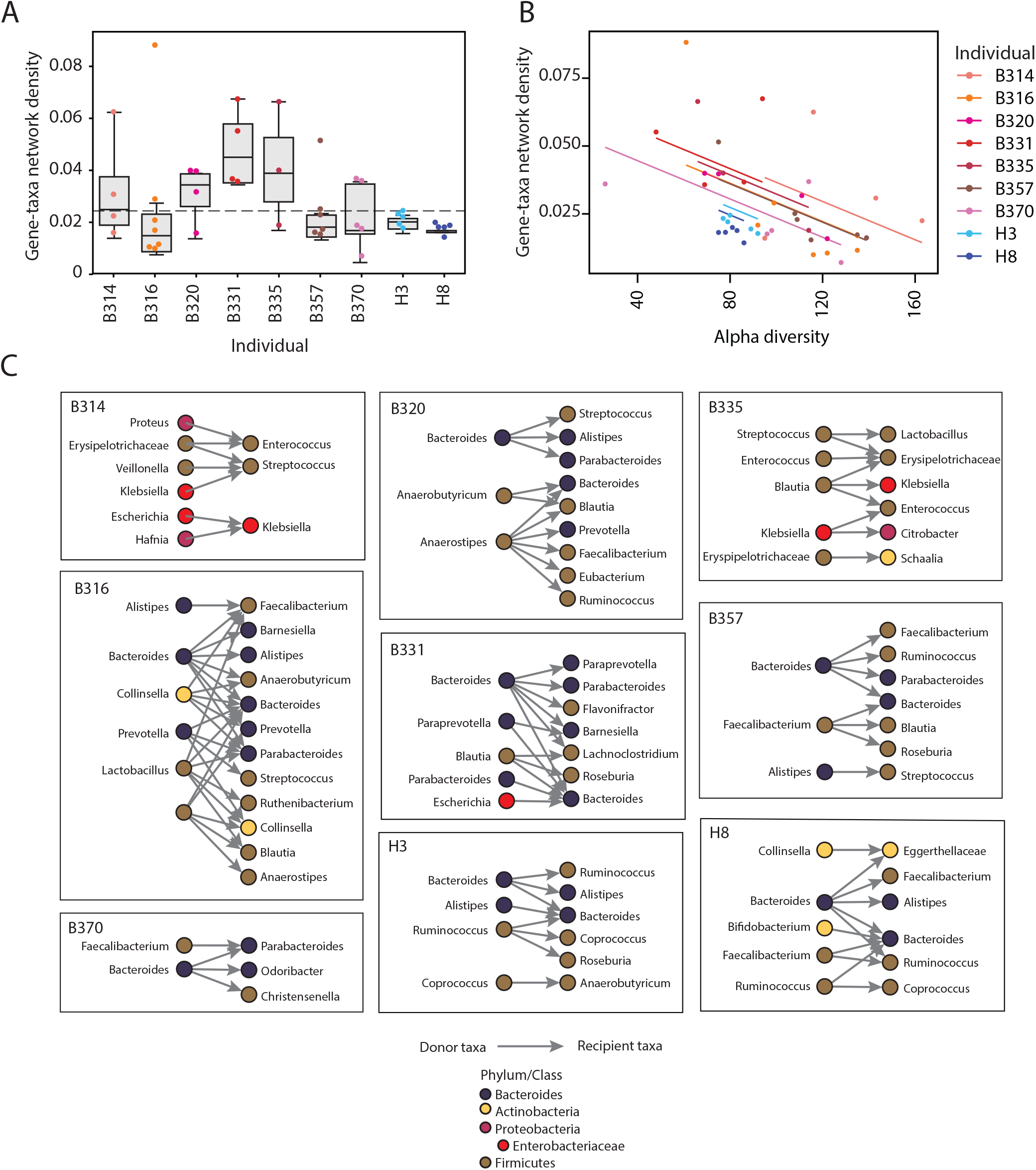
HGT results in greater network density and occurs on short timescales in some members of our neutropenic cohort compared to healthy individuals. (A) Boxplots showing the gene-taxa network linkage densities, or the proportion of total possible gene-taxa links that are observed to be linked, for each individuals’ samples. A dotted line is shown at the maximum network density observed in the healthy samples. (B) Individual patient samples are plotted according to the alpha diversity, assessed using Metaphlan, and their gene-taxa network density. An ANOVA showed that gene-taxa network density was related to alpha diversity (F(1,39), p = 3.3×10^−5^) and health status (F(1,6), p=0.01501). (C) All observed HGT events across different genera are plotted for each individual. Each genus is colored according to its phylum.

Next, we more closely examined those timecourses with putative HGT events for which we had the highest confidence. To distinguish between the migration of new bacterial strains and HGT, we only considered HGT between strains present at the start of the timecourse. Potential donor strains were required to have Hi-C-verified connections with specific mobile or AR genes in the first patient sample. Individuals’ initial Hi-C samples were sequenced 3-4-fold deeper than the remainder of their timecourses to ensure that gene-taxa associations were adequately sampled (Extended Data Figure 9) and that putative recipient strains did not harbor those specific genes of interest at the start. We enforced this by requiring a complete absence of gene-recipient taxa connections inferred by Hi-C or metagenomic assembly, including connections with taxa that could only be annotated at higher taxonomic levels. Finally, we considered HGT as occurring between these donor strains and recipient strains with Hi-C-verified associations with the transferred genes in later timepoints. Providing additional support, 12.2% of the putative transfer events (19 of 155) were supported by Hi-C across multiple timepoints and 32.9% (51 out of 155) were supported by Hi-C links across multiple contigs in both the donor and recipient genomes. Most of the transfers (60%) were between members of the same phylum. Ultimately, evidence of HGT was found in all individuals in our study (Figure 3C, Extended Data Table 6).

Within these relatively short timecourses, we observed the expansion of the gut commensal reservoir of resistance genes. Although we did not observed the transfer of AR genes conferring resistance specifically to the antibiotics used in this cohort, namely levofloxacin, pipercillin-tazobactam, or trimethoprim-sulfamethoxazole, we did observe transfer of multi-drug resistance cassettes with beta-lactam- and fluoroquinolone-resistance genes, covering two of the corresponding antibiotic classes. Notably, within a few days post-transplant, we see transfer of a plasmid encoding mdtEF, a multidrug efflux pump conferring resistance to fluoroquinolones, and their transcriptional regulators, CRP and gadW, from an *Escherichia coli* strain in patient B331 to a strain most similar to *Bacteroides sp. A1C1*. Despite their ubiquitous antibiotic prophylaxis, only a minority (19.4%) of transfer events involved annotated AR genes in the neutropenic patients.

Additionally, we observe the emergence of novel AR genes in enteric pathogens, originating either from gut commensals or other enteric pathogens, including Enterobacteriaceae. Enterobacteriaceae species are among the most common causes of infection and sepsis in these patients and Enterobacteriaceae from the gut have been shown previously to harbor excessive numbers of AR genes^41^ and serve to promote HGT of AR genes^42^. We see the exchange of AR gene-containing plasmids between members of the Enterobacteriaceae, namely *Klebsiella pneumoniae* and *Citrobacter brakii* in patient B335, and between *E. coli* and *Klebsiella* species in B314, and one instance of *K. pneumoniae* in patient B335 acquiring DNA harboring a plasmid-based efflux pump from a commensal, *Blautia hansenii*. We also note the overall transfer of mobile elements between these pathogenic species and other opportunistic pathogens, such as *Streptoccocus parasanguinis*, *S. salivarius*, and *E. faecium*, exposing the potential for HGT to alter the AR profiles of these bacteria over short periods of time.

These examples highlight the dynamic nature of HGT within the gut ecosystem, especially in the context of gut inflammation, immune dysregulation and antibiotic use. Nevertheless, our method has several limitations. First, we can only assign bacterial hosts for those MGEs and host genomes that we are able to assemble and annotate. Although 95.9% ± 2.8% of our metagenomic reads contribute to assembled contigs and 80.4% ± 10.4% (median 82.4%) of Hi-C reads align to our assemblies, we were only able to annotate 47.8% of our draft assemblies at either the genus- or species-level. Second, the assembly of MGEs can be confounded by their high rates of recombination creating multiple genomic arrangements and transfer, and, thus, redundancy within and across genomes. To mitigate the potential for false positive interactions, we examined only those mobile gene-containing contigs with multiple Hi-C reads directly linking them to taxonomically annotated genome assemblies. We cannot however rule out the possibility that our sensitivity is actually higher, and that our inability to detect linkages at specific time-points reflects true strain-level variation within the microbiome, or undetected real-time mobilization of genetic elements. Third, for those HGT events that we observed, we cannot always confirm the transfer of an entire contig and its associated genes. This issue underscores several observed HGT events, involving plasmids comprising prophage and transposable elements. This is mitigated by the requirement for more Hi-C read linkages and the overall proximity between Hi-C read linkages and the inferred transferred genes (Extended Data Figure 7). Future studies should leverage long reads in hybrid assemblers to better capture co-occurring AR genes and large MGEs^43^. We expect to overcome these limitations with additional technical improvements to the bacterial Hi-C protocol.

Here, we observe extensive transfer of mobile and AR genes within a single individual’s gut microbiome across distant phylogenetic backgrounds and over relatively short timespans. The transfer networks within each individual’s gut microbiome are unique and are likely explained by personal ecological niches that govern local contact rates between organisms. Few of the total AR gene-taxon associations are observed across individuals, which may suggest limited dispersal rates and/or strong selective pressures that prevail within each individual’s gut. Although the molecular dynamics of HGT in the gut microbiome are not well-understood, our data from healthy subjects point to a basal level of transmission, even in the absence of inflammation or antibiotic use. Many of the observed transfers appeared transient, which may be due to the limited detection of our method, or by the neutral or deleterious nature of most HGT events^44,45.^

The ramifications of HGT in this neutropenic patient population are acute. Our results show increased pathogen load and elevated gene-taxa network densities in neutropenic patients as compared with healthy individuals, suggesting an increased risk of emergence of MDR pathogens in this at-risk patient population. How to translate these findings into the prevention of the emergence of MDR pathogens is paramount. This technology highlights the potential for screening the burden of AR genes and the carriage of enteric pathogens to guide empirical antibiotic therapy. These findings also expose the limitations of taxa-specific therapies to remove AR genes from the gut microbiome^46,47,48^, in favor of mechanisms to limit HGT more generally. Overall, these results emphasize a view of the population-wide dissemination of AR genes that includes diverse members of the gut microbiome.

## Extended Data Figures

**Extended Data Figure 1.**
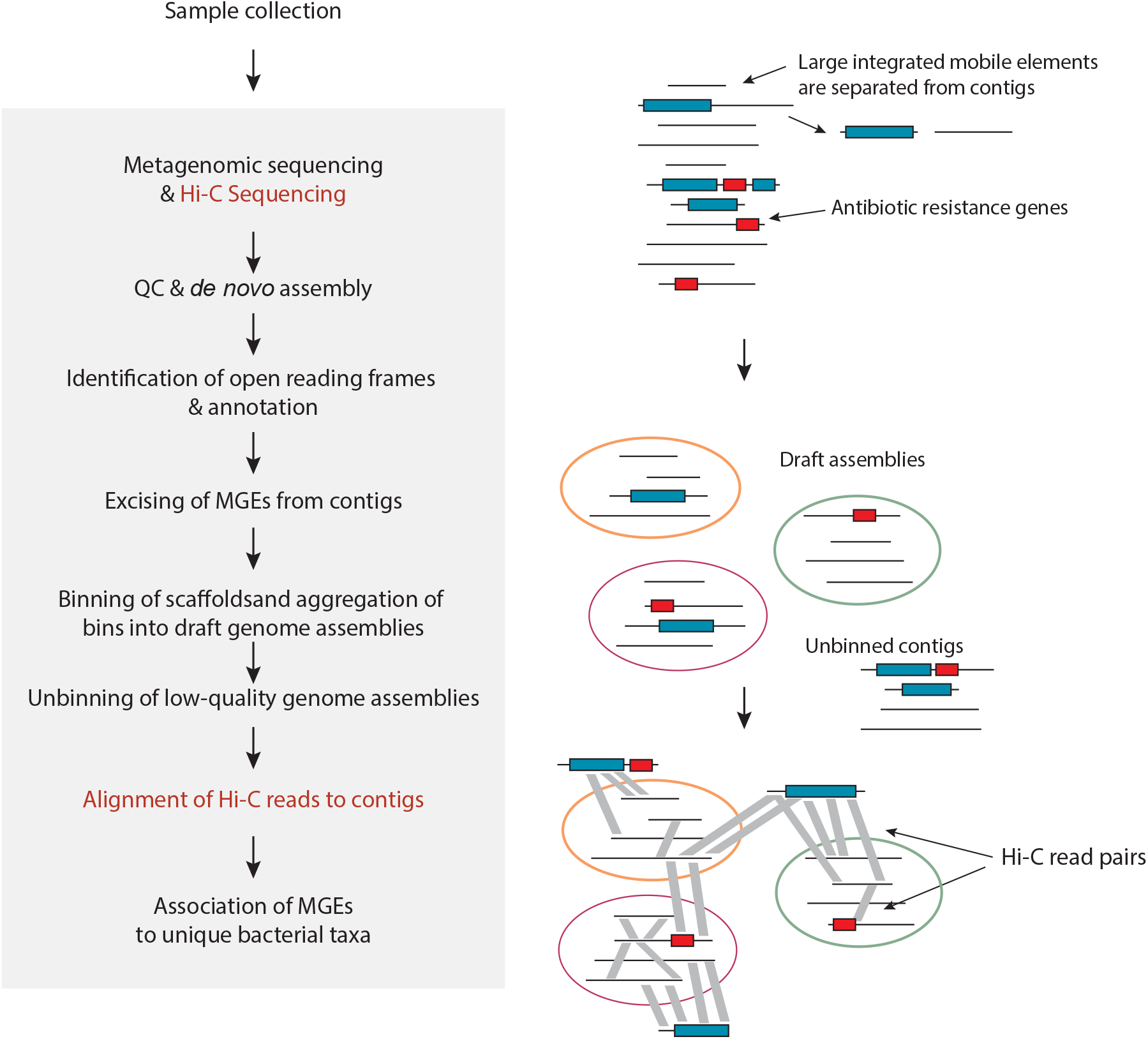
Experimental and computational pipeline. Our pipeline for assigning mobile and AR genes to bacterial taxa utilizes data from metagenomic (black) and Hi-C (red) sequencing libraries. In brief, metagenomic samples are assembled using standard approaches. The resulting contigs (circles) are binned into draft assemblies and quality filtered. Bins are taxonomically annotated at the lowest level with >50% of bps assigned to a taxon using a weighted Kraken approach. Contigs containing AR or mobile genes are associated with metagenomic assemblies by residency or by Hi-C linkages requiring at least 2 readpairs linking the contig with the metagenomic assemblies. Associations of mobile or AR genes with specific taxa are made by clustering the genes of interest at 99% identity and counting each unique taxon once.

**Extended Data Figure 2.**
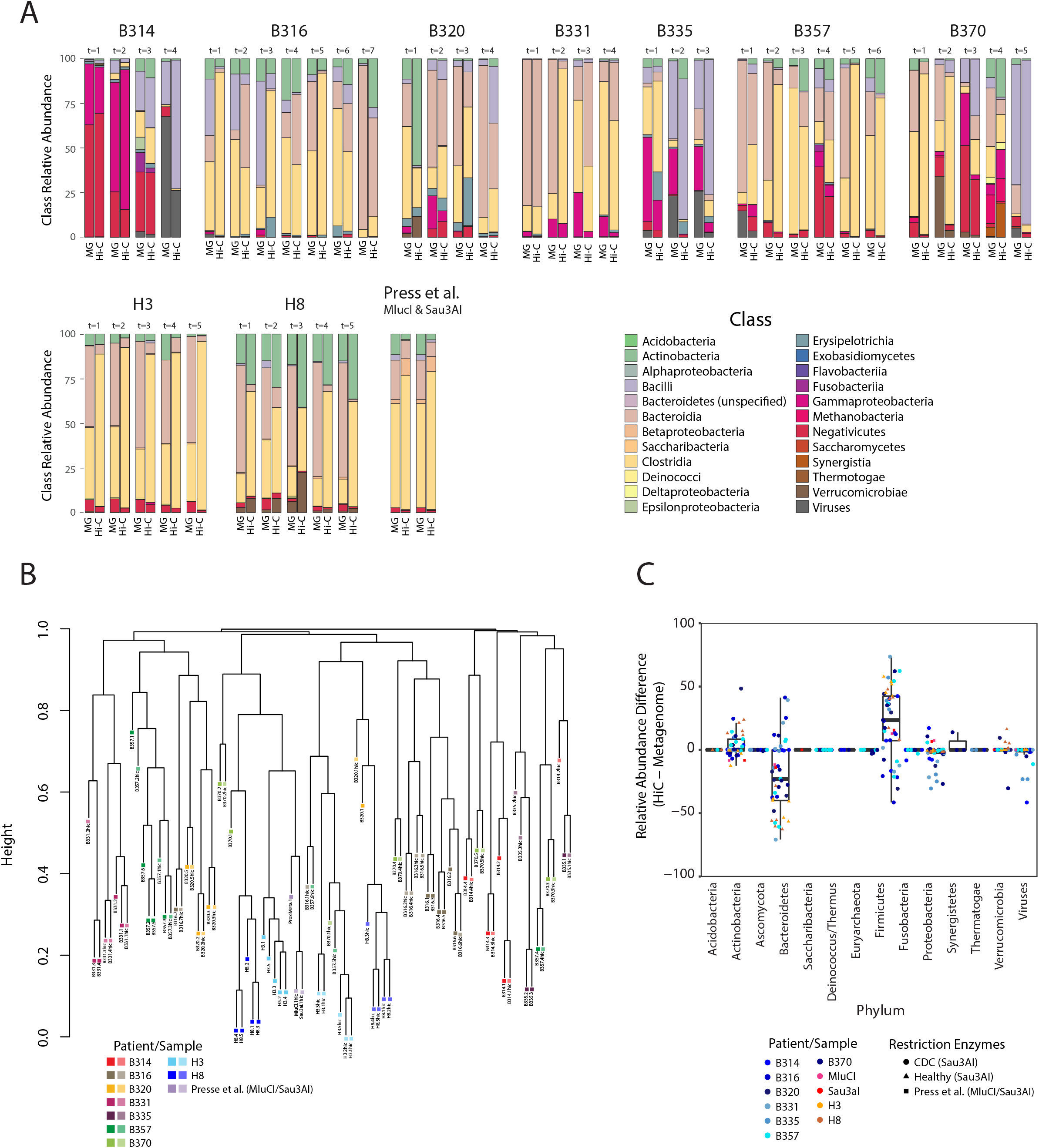
Congruence between metagenomic and Hi-C sample composition. (A) Class-level compositions of individuals’ gut metagenomes and Hi-C libraries as determined by MetaPhlAn2. The human microbiome sample from Press *et al^49^*. was included in our analysis. (B) Dendrogram of metagenome and Hi-C library compositions. Sample compositions (class-level) were hierarchically clustered according to their Bray-Curtis distances. (C) Compositional differences between metagenomic and Hi-C libraries in samples processed according to the restriction enzyme(s) used.

**Extended Data Figure 3.**
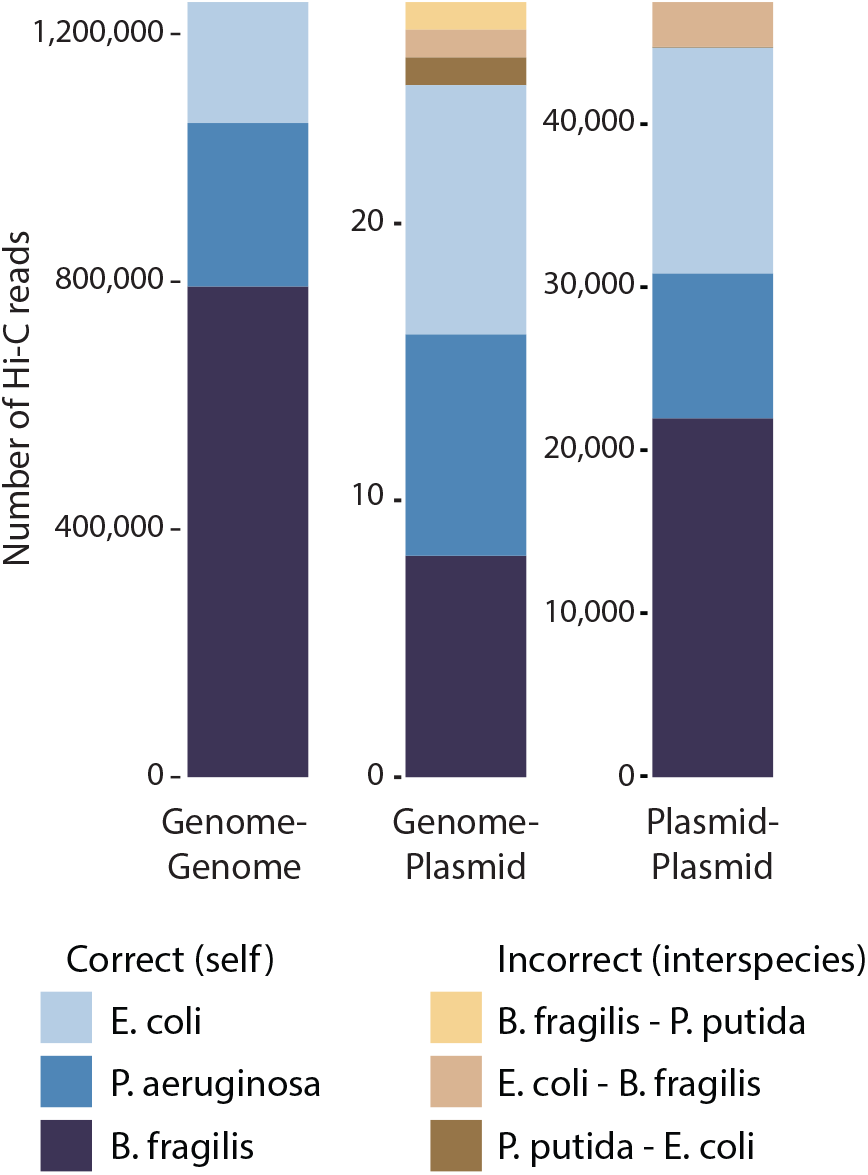
Hi-C performed on a mock community of three organisms each harboring an organism-specific plasmid. Number of Hi-C reads linking genomic regions to themselves (left), to their plasmids (middle) and within each plasmid (right). Blue linkages are correct, whereas brown hues are incorrect associations. Note that there is a region of homology between the plasmid backbones of the RP5 and pKJK5 plasmids carried by *Pseudomonas putida* and *Escherichia coli*, respectively. Nevertheless, none of the incorrect host-plasmid linkages would have been surpassed our threshold for assigning gene-taxa associations.

**Extended Data Figure 4.**
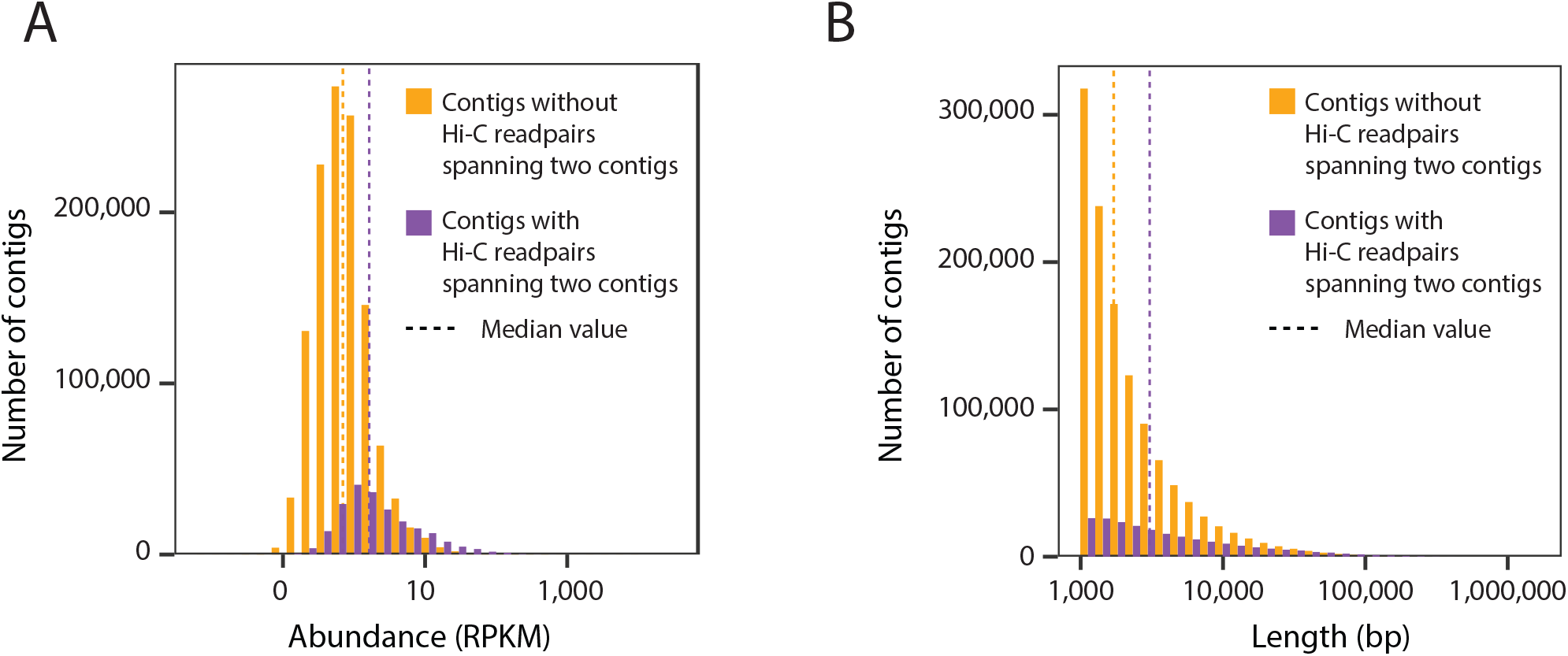
Recruitment of Hi-C read pairs linking separate contigs according to the length and abundance of those contigs. (A) A histogram showing the distribution of abundances (RPKM) of contigs that recruit (purple) or do not recruit (orange) Hi-C contig-connecting read pairs. (B) A histogram showing the distribution of lengths (bp) of contigs that recruit (purple) or do not recruit (orange) Hi-C contig-connecting read pairs.

**Extended Data Figure 5.**
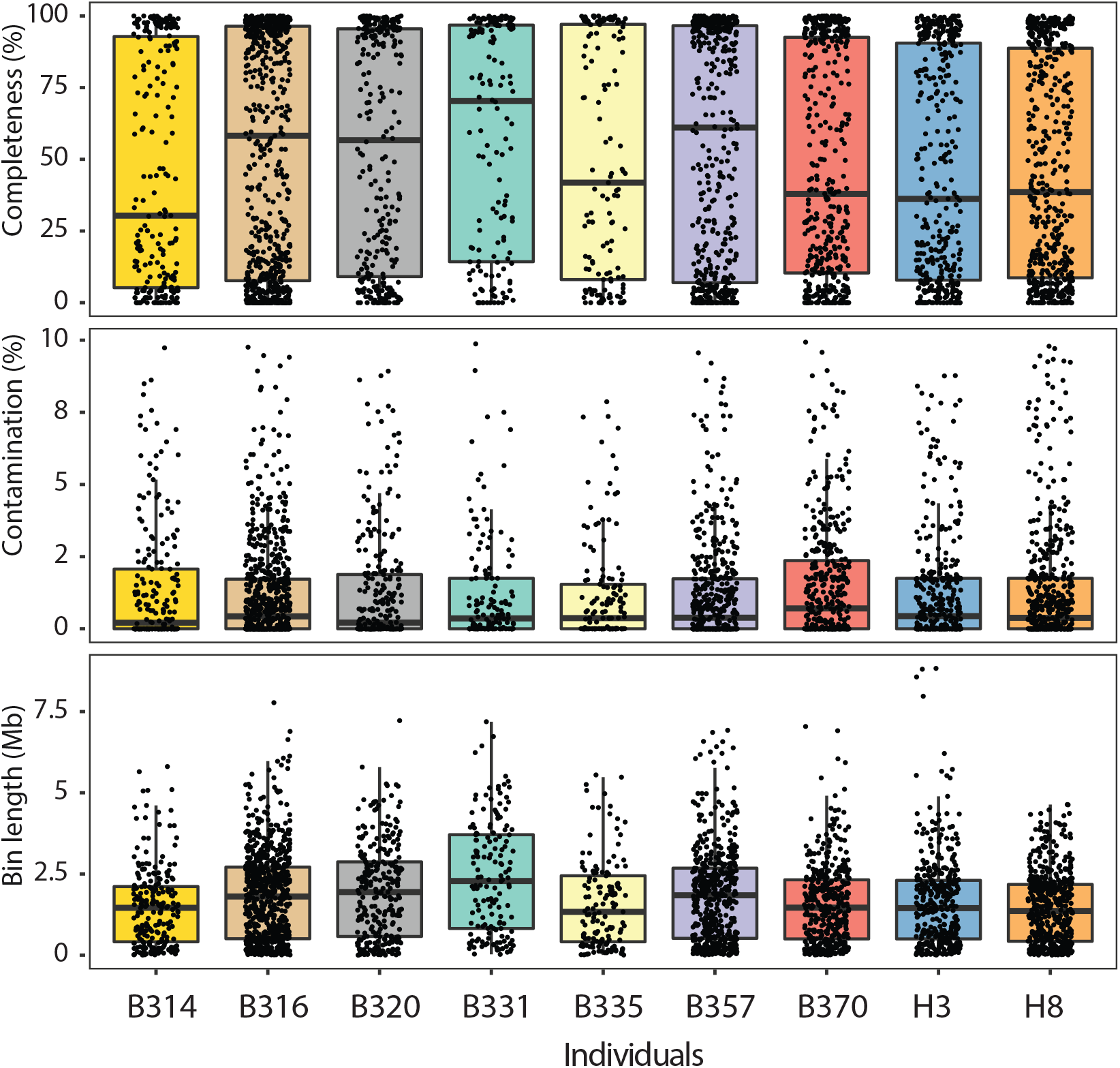
Genome bins are high-quality in terms of completeness and contamination. (A) Completeness of each assembled genome bins from each patients’ samples as scored by CheckM. Boxplots show 25^th^, median and 75^th^ percentile. (B) Contamination of each assembled genome bins from each patients’ samples as scored by CheckM. Boxplots show 25^th^, median and 75^th^ percentile. (C) Length of each assembled genome bins within each patients’ samples. Boxplots show 25^th^, median and 75^th^ percentile.

**Extended Data Figure 6.**
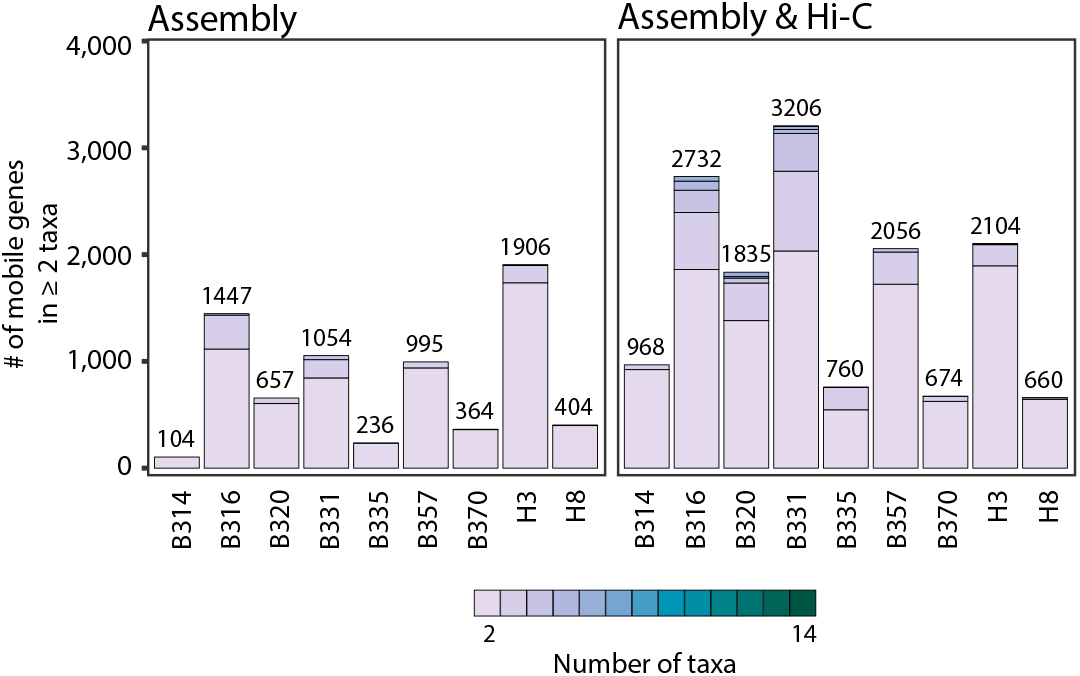
Hi-C associates mobile genes with multiple taxa. Stacked barplots showing the number of species-level taxa to which each mobile gene (clustered at 99% identity) is assigned within each patient, and across patients. Only those genes assigned to 2 or more taxa are shown. We either used metagenomic assemblies alone to assign taxonomies (left) or Hi-C libraries considering those taxa-gene assignments with evidence from at least two Hi-C read pairs. The numbers above each stacked barplot represent the total number of mobile genes with 2 or more taxonomic associations.

**Extended Data Figure 7.**
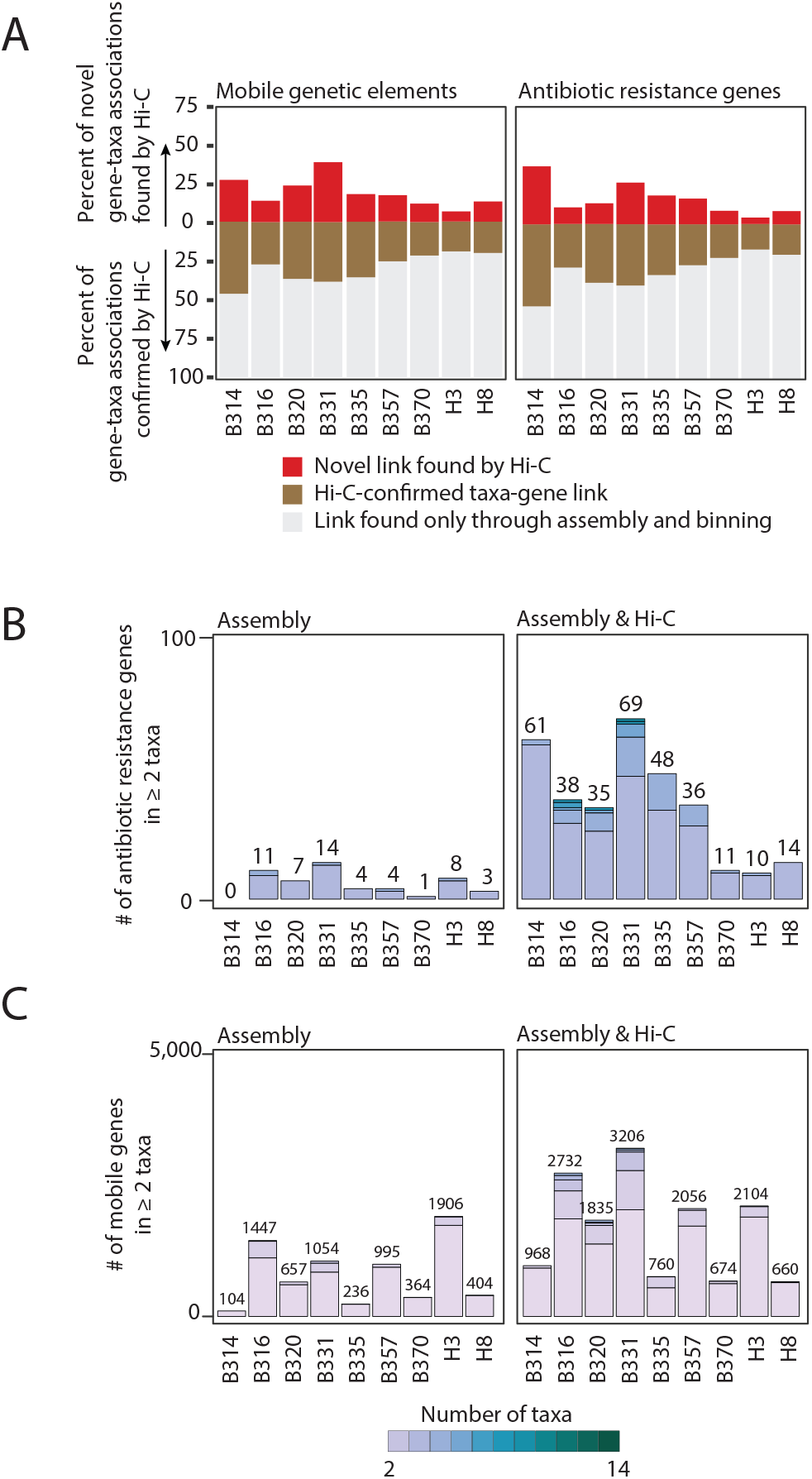
Trends in gene-taxa associations are consistent with more stringent cutoffs of Hi-C read linkages. (A) The percent of the total taxa-mobile gene (left) and taxa-AR gene (right) associations observed from metagenomic assembly that are supported by five or more Hi-C links (brown) is plotted, along with the percent additional interactions gained by using Hi-C (red). (B) Stacked barplots showing the number of species-level taxa to which each AR gene (clustered at 99% identity) is assigned within each patient, and across patients as determined by 5 or more Hi-C links. Only those genes assigned to 2 or more taxa are shown. We either used metagenomic assemblies alone to assign taxonomies (left) or combined with Hi-C libraries considering those taxa-gene assignments with evidence from at least two Hi-C reads. The numbers above each stacked barplot represent the total number of AR genes with 2 or more taxonomic associations. (C) Stacked barplots showing the number of species-level taxa to which each mobile gene (clustered at 99% identity) is assigned within each patient, and across patients as determined by 5 or more Hi-C links. Only those genes assigned to 2 or more taxa are shown. We either used metagenomic assemblies alone to assign taxonomies (left) or combined with Hi-C libraries considering those taxa-gene assignments with evidence from at least two Hi-C reads. The numbers above each stacked barplot represent the total number of AR genes with 2 or more taxonomic associations.

**Extended Data Figure 8.**
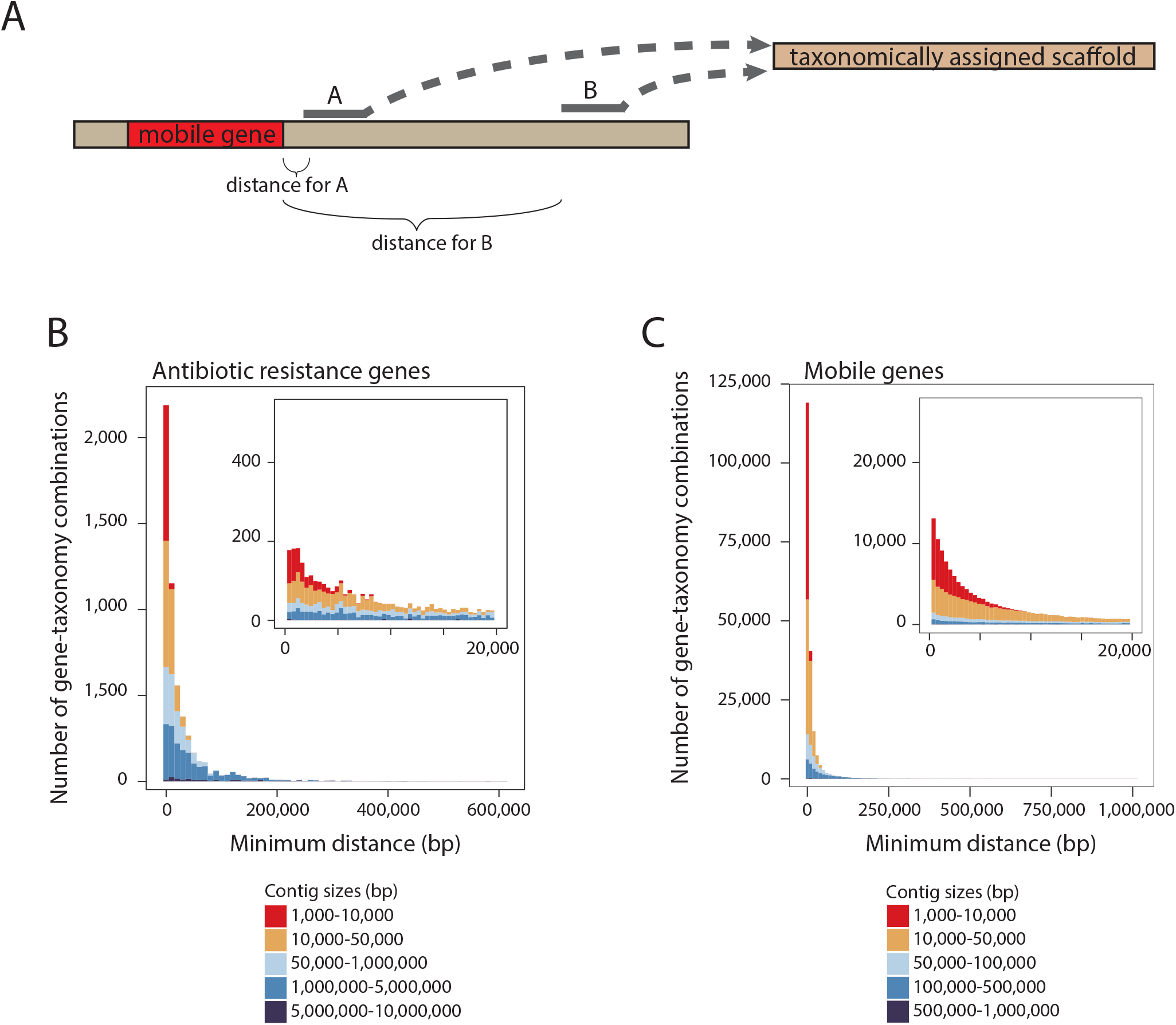
Hi-C read pairs are situated nearby mobile or AR genes on contigs. (A) As illustrated, Hi-C read pairs may map anywhere on a contig containing a mobile or AR gene. The boundaries of an MGE may be elusive and MGEs may integrate into contigs that have incomplete annotations. We assessed the linear distance (bp) between where Hi-C read pairs aligned and the positions of mobile or AR genes used for taxon-gene associations on the contigs to ensure that Hi-C read pairs were mapping at distances relevant for their assignments. In the example, both Hi-C reads A and B align to the same taxonomically annotated contig, yet read A maps at a minimum distance that is closer to the mobile gene, and therefore more confidently links the mobile gene with the contig. (B) For each taxon-mobile gene connection, we plot the minimum distance between a Hi-C read pair and the start/end of the mobile gene on that contig, according to contig length. The inset shows distances of less than 200,000bp broken down more finely. (C) The same analysis as (B) but for AR genes.

**Extended Data Figure 9.**
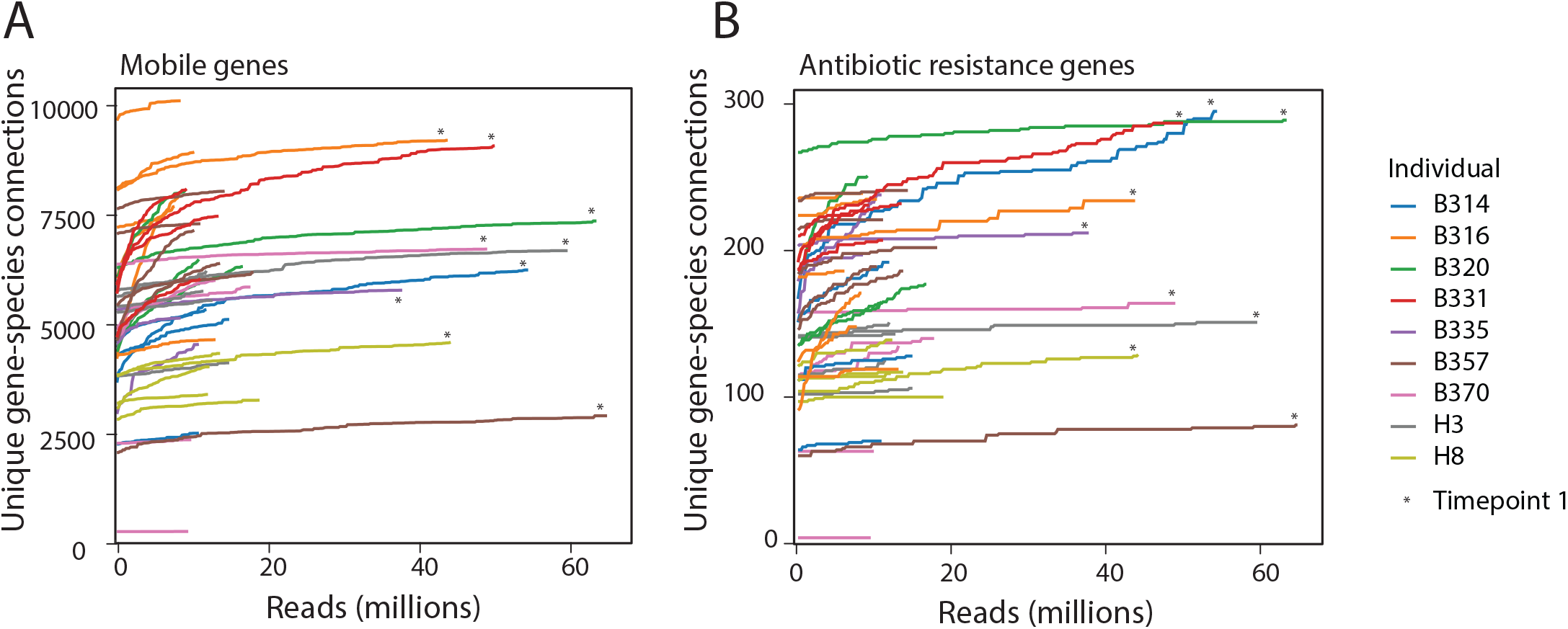
Accumulation of unique gene-species connections with increasing sequencing depth. (A) The number of unique mobile gene-species connections within each sample after subsampling reads from each Hi-C dataset. The first Hi-C sample from each timecourse (noted with an asterisk) was sequenced significantly more deeply than the rest of the timecourse so that we could better assess whether there were any new gene-species connections that arose in any subsequent samples. (B) The number of unique AR gene-species connections within each sample after subsampling reads from each Hi-C dataset.

**Extended Data Figure 10.**
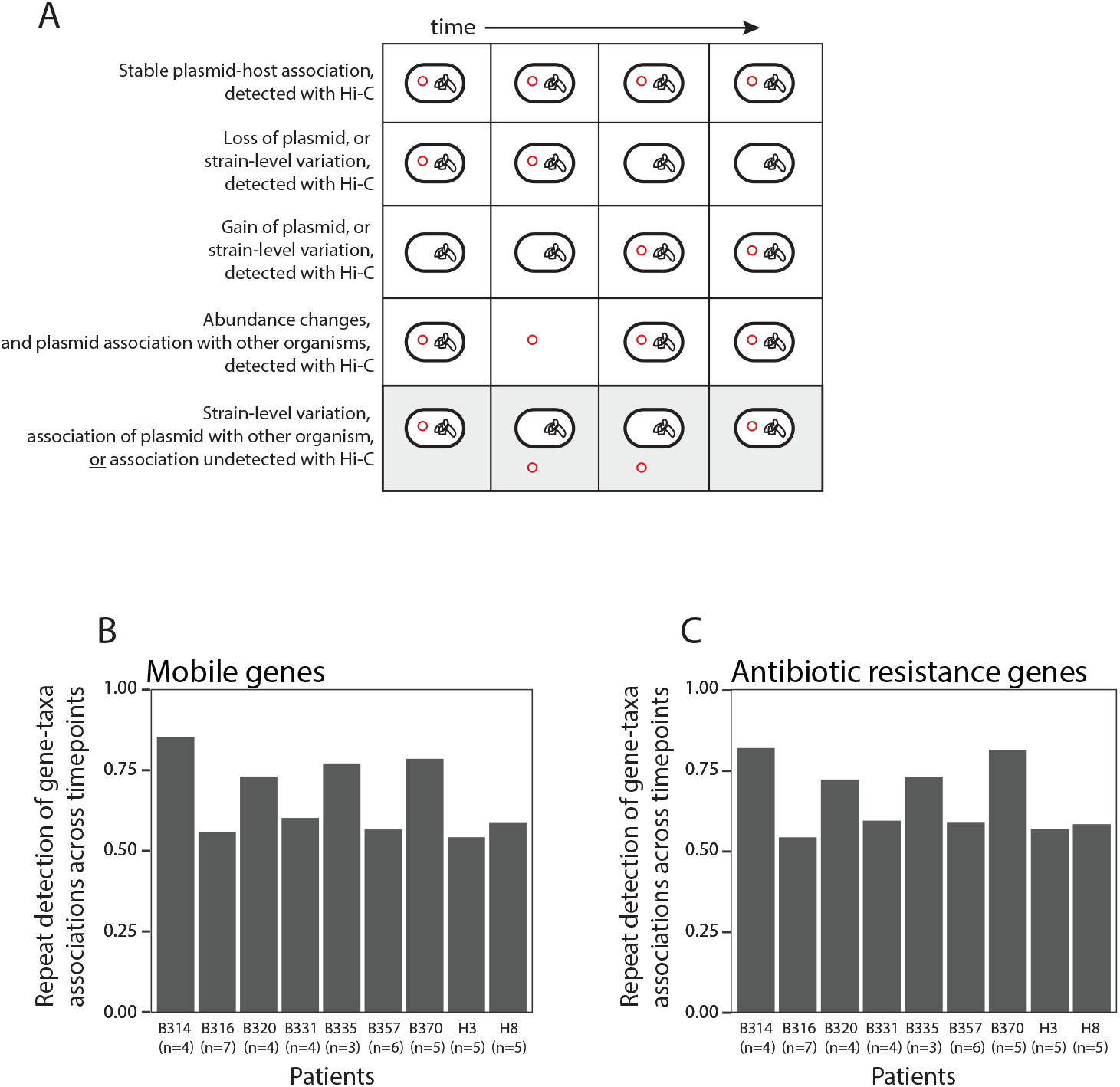
Repeat detection of gene-taxa associations within the Hi-C libraries. (A) Examples of the detection of AR genes, bacterial hosts and their linkage during patients’ timecourses. The last scenario depicts an instance where a mobile or AR gene is linked with a specific taxon with Hi-C during at least one time point, but is detected in the metagenomic data at other time points but not linked with Hi-C. Although this may be explained by changes in strain-level composition or gene loss, we assessed the repeatability of detecting associations, assuming that these genes are truly linked in any instance when the mobile or AR gene and the bacterial taxon are both present. (B) A bar chart showing the extent to which we repeatedly detect specific gene-taxa associations observed within each patients’ microbiomes. Assuming that we should observe associations present in one timepoint in all timepoints (*i.e.* that there is no HGT), we define the true positives (TPs) as the number of unique mobile gene-bacterial taxon connections observed; and the false negatives (FNs) as the total number of instances where both the bacterial taxon and the mobile gene are detected in the metagenomic assemblies. Repeat detection is calculated as TPs/(TPs+FNs), with the caveat that a portion of genome mobile gene-taxon linkages that did not depend on Hi-C sequencing read pairs are included here. (C) A bar chart showing the amount of repeat detection of AR gene-taxon connections observed within each patients’ microbiomes. This was calculated as described in (B), with the same caveat applied.

**Extended Data Figure 11.**
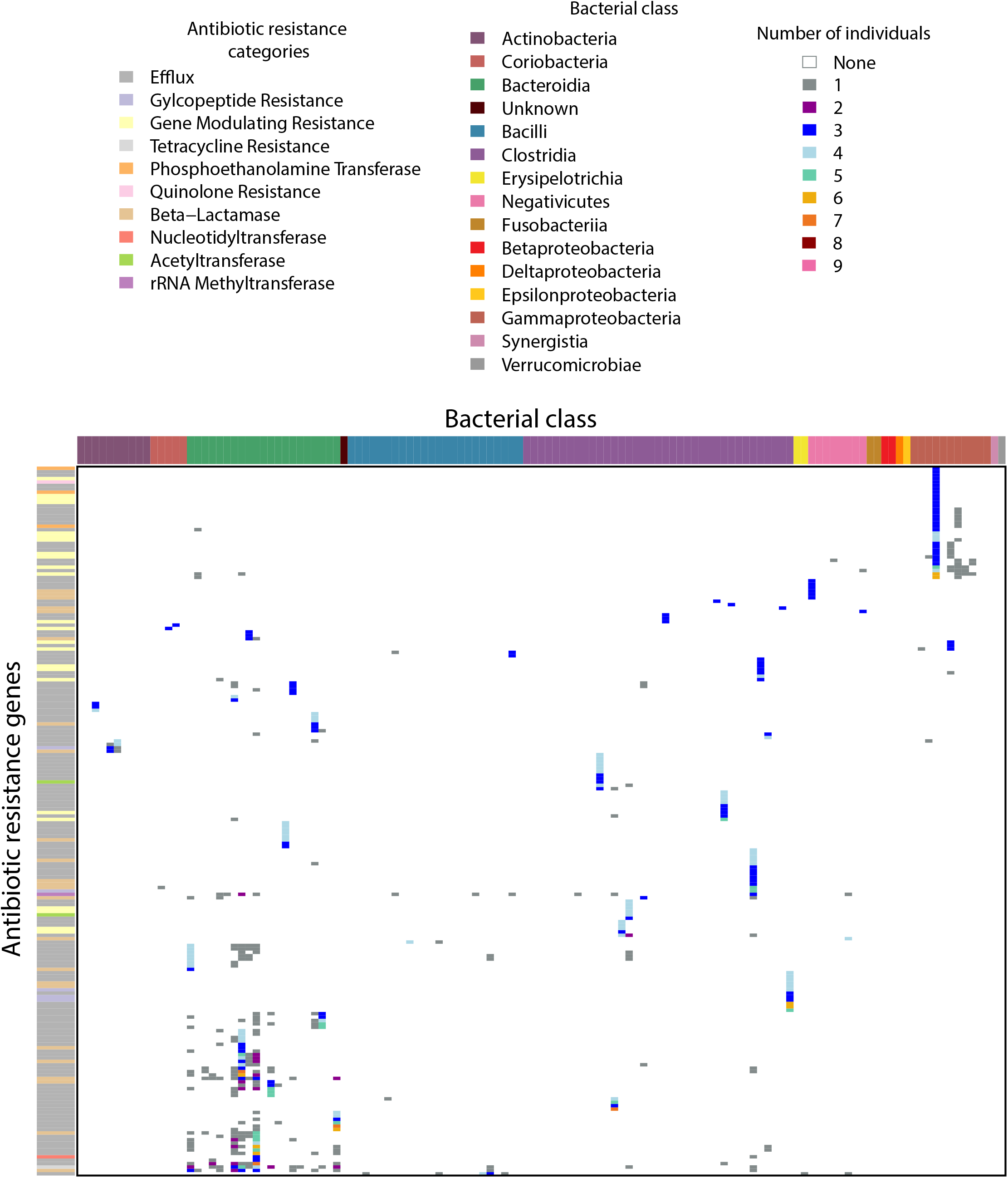
AR gene-taxa linkages are not common across individuals. A heatmap of taxa-specific assignments, colored by class, for AR genes that are present in three or more patients.

**Extended Data Figure 12.**
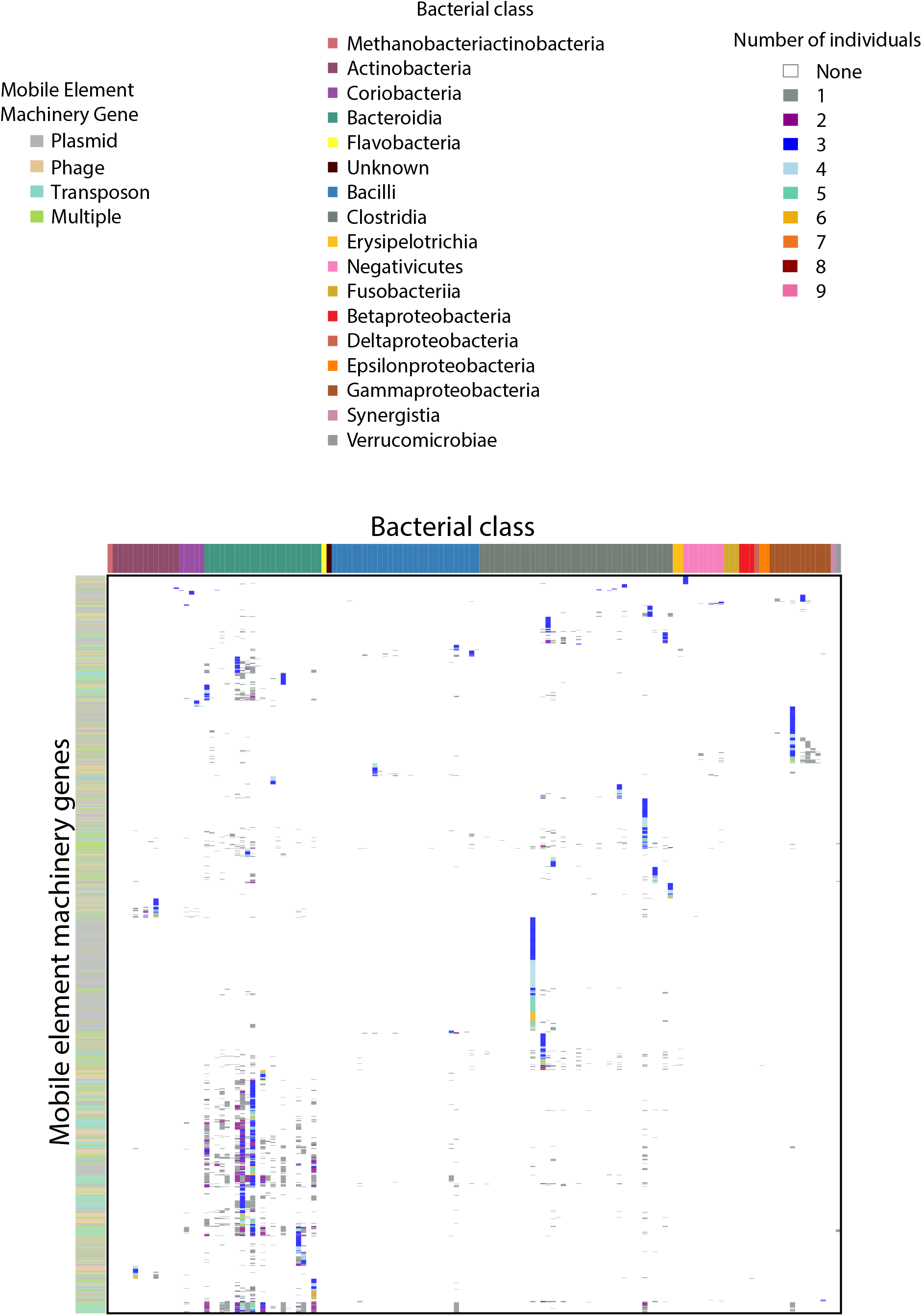
Mobile gene-taxa linkages are not common across individuals. A heatmap of taxa-specific assignments, colored by class, for mobile genes that are present in three or more patients.

**Extended Data Figure 13.**
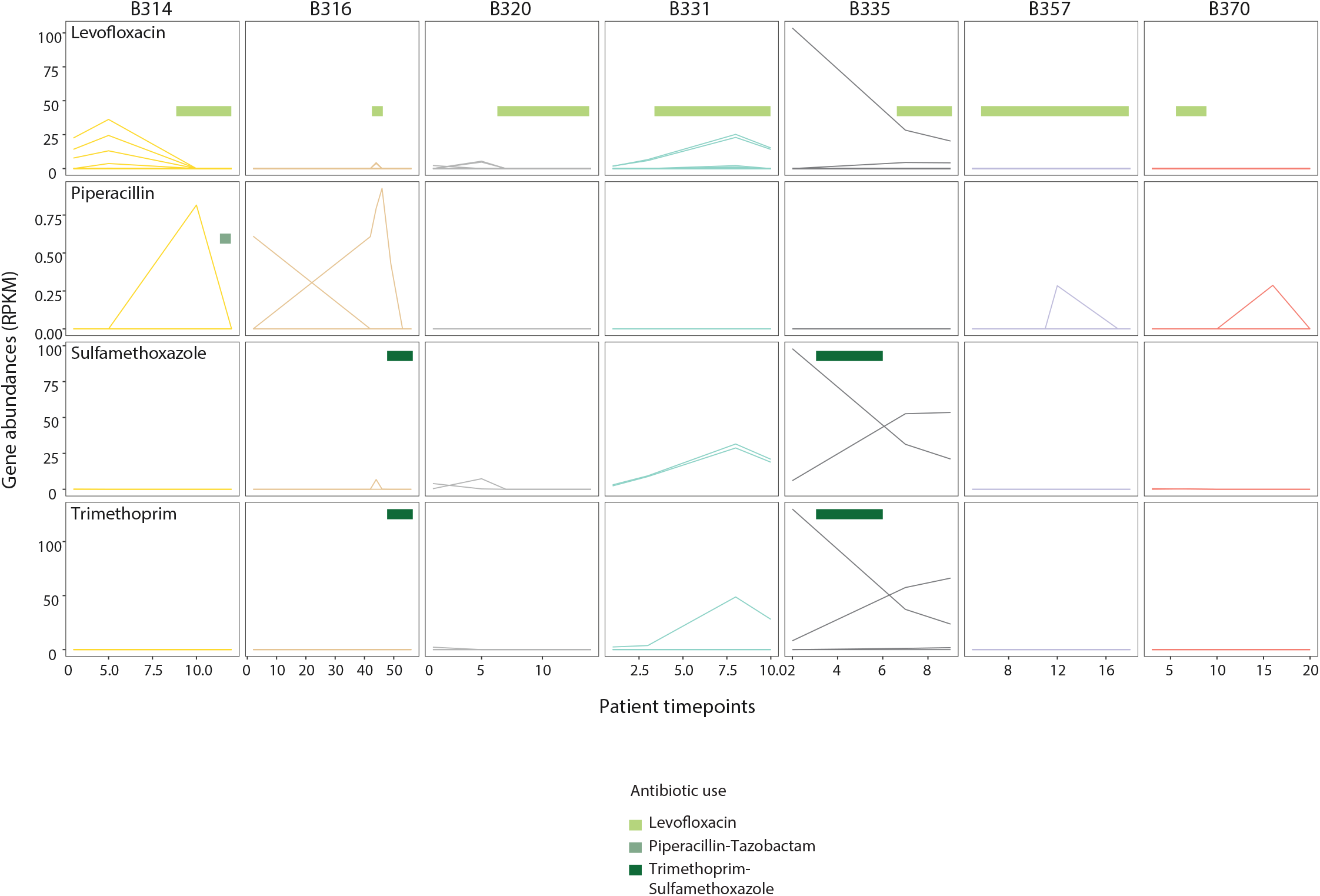
Antibiotic resistance gene abundances do not correspond to patients’ antibiotic regimens. For each patient-timecourse (columns), AR gene abundances (RPKM) are plotted for each gene according to the antibiotic to which it confers resistant. The antibiotics administered to each patient over their timecourse is denoted.

**Extended Data Figure 14.**
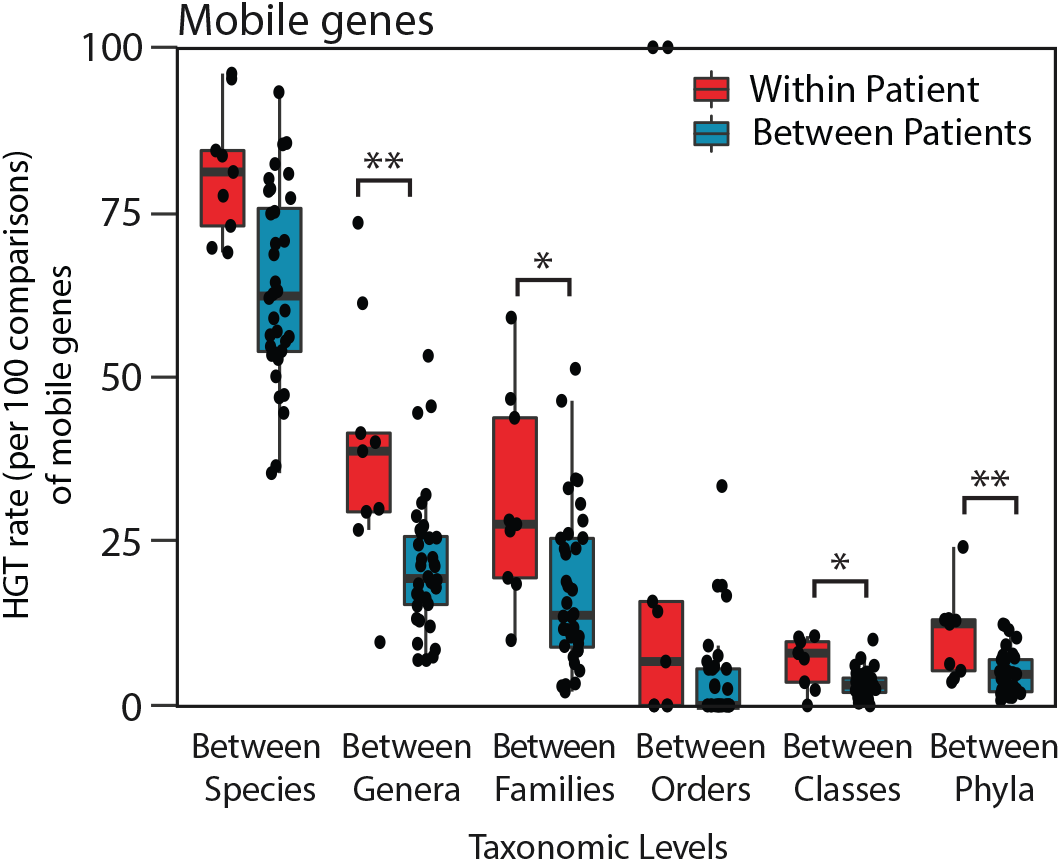
Transfer of AR genes between organisms is higher within individuals than across individuals. HGT rates (per 100 comparisons) of AR genes between organisms within each individual versus between individuals, according to those that share the same genus, family, order, class, phylum and kingdom are plotted for comparison. Significance was measured with Mann Whitney U-tests (two-sided; *, p<0.05; **, p<0.01; ***, p<0.005, ****, p<0.001;*****, p<0.0005).

**Extended Data Figure 15.**
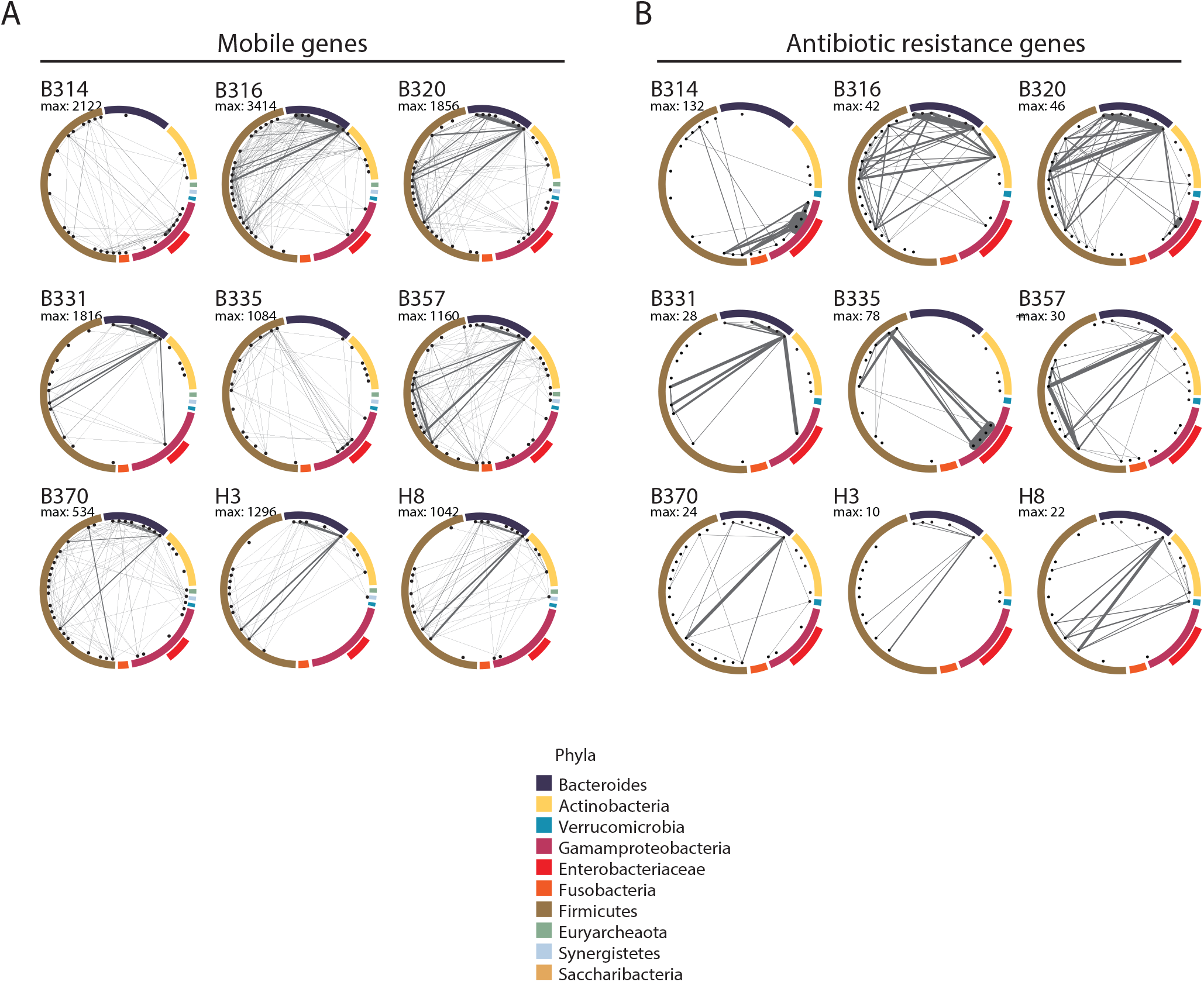
Networks of bacterial HGT of AR genes are unique to each individuals’ gut microbiomes. (A) Circle relationship plots showing networks of bacterial mobile gene exchange in the gut microbiomes across individuals (top left) and within each individual. Taxa present in each individual’s microbiome are depicted by black circles. The thickness of the lines corresponds to the number of unique mobile genes associating the two taxa. (B) Circle relationship plots showing networks of bacterial AR gene exchange in the gut microbiomes across individuals (top left) and within each individual. Taxa present in each individual’s microbiome are depicted by black circles. The thickness of the lines corresponds to the number of unique AR genes associating the two taxa.

**Extended Data Figure 16.**
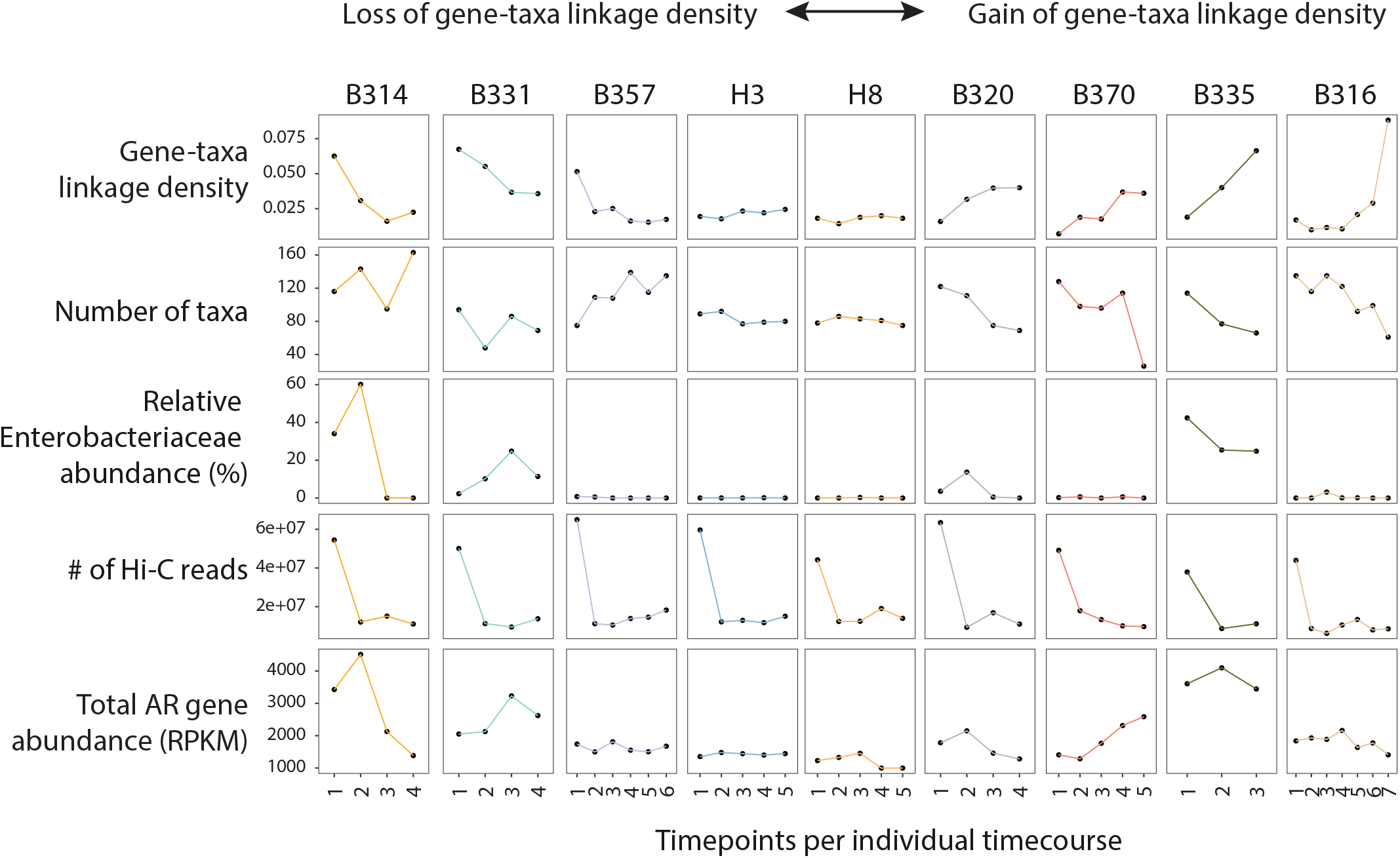
Gene-taxa linkage density is related to the number of taxa, but not to the relative abundance of Enterobacteriaceae, number of Hi-C reads or abundance of AR genes. Each patient’s timecourse is shown according to their gene-taxa linkage density, as defined by both Hi-C and metagenomic assembly; the number of taxa in that sample’s metagenome (calculated by Metaphlan), the number of total Hi-C reads; and the abundance of AR genes (RPKM). Individuals were ordered according to the trend of their gene-taxa linkage density over their timecourse.

## Extended Data Tables

**Extended Data Table 1. Patients and clinical data**

Patient data, including type of stem cell transplantation, dates of admission, pre-transplant conditioning (radiation, chemotherapy, or mAb therapy), neutropenia, and antibiotic treatment are provided, relative to the date of hematopoietic cell transplantation.

**Extended Data Table 2. Sample quality and information**

For each patient-time point sample, quality information is provided about the metagenomic and Hi-C libraries. Metrics included are the read depth and assembly metrics of the metagenomes, the number of Hi-C reads, restriction enzymes used in the Hi-C data, percent of inter-contig chimeras.

**Extended Data Table 3. Completeness and contamination of genomic clusters.**

We clustered contigs using Maxbin, MetaBat, Concoct, and DAS. We ran CheckM on all of the clusters and report completeness, heterogeneity, and contamination. We applied a bp weighted taxonomic annotation of Kraken contig assignments for each bin and annotated lowest taxonomic levels that attained >50% of bin length.

**Extended Data Table 4. Antibiotic resistance mechanisms**

AR genes were identified using HMMer against the Resfams database, and using CARD’s Resistance Gene Identifier against the CARD database. Due to slight variations in AR gene classification, we combined annotations according to the mechanism specified in this look-up table. We used a slightly finer resolution categorization for the subset of clinically relevant genes.

**Extended Data Table 5. Mobile PFAMs included in this study.**

A table of the PFAM IDs that were used, according to search terms described in the Methods, and how they were annotated according to MGE (*i.e.* plasmid, phage, transposon, mixed).

**Extended Data Table 6. Putative HGT events during individuals’ timecourses.**

For each patient, we list the observed HGT events that meet the inclusion criteria outlined in the Methods. We include the full taxonomies of the donor and recipient taxa, the gene IDs and gene names involved in the transfer, whether specific genes were found to be transferred across multiple timepoints, whether there was any evidence of transfer between two taxa across multiple timepoints, and whether multiple contigs were associated with transfer between two taxa at a single timepoint.

## Methods

### Sample collection

Fresh stool was collected from individuals in accordance with IACUC protocol for Weill Cornell Medical College (#1504016114) and Cornell University (#1609006586). Neutropenic patients were all admitted to the Bone Marrow Transplantation Unit at New York Presbyterian Hospital/Weill Cornell Medicine between December 2016 and July 2017. Healthy samples were collected similarly in 2019. Approximately 0.25 g replicates of each time-point were either frozen ‘as is’ (for metagenomic sequencing) or homogenized in phosphate-buffered saline (PBS) + 20% glycerol before freezing (used for Hi-C sequencing).

### Metagenomic sequencing

Frozen stool was thawed on ice and DNA was extracted using the PowerSoil DNA Isolation Kit (Qiagen) with additional Proteinase K treatment and freeze/thaw cycles recommended by the manufacturer for difficult-to-lyse cells. Extractions were further purified using 1.8 volumes of Agencourt AMPure XP bead solution (Beckman Coulter). DNA was diluted to 0.2 ng/uL in nuclease-free water and processed for sequencing using the Nextera XT DNA Library Prep Kit (Illumina).

### Proximity Ligation

Stool stored in PBS + 20% glycerol was thawed on ice for 15 minutes and homogenized in 5 mL PBS containing 4% v/v formaldehyde. Sample were crosslinked at room temperature with continuous inversion for 30 minutes, then incubated on ice for 30 minutes. Unreacted formaldehyde was quenched by adding glycine to a final concentration of 0.15 M and incubating for 10 minutes on ice. Crosslinked cell mixtures were pelleted (10,000 g, 4° C, 5 min.), the supernatant was removed, and pellets were flash-frozen on dry ice/ethanol and stored at −80° C.

Frozen crosslinked stool cell pellets were thawed on ice then resuspended in 450 μL TES (10 mM Tris, 1 mM EDTA, 100 mM NaCl, pH 7.5) and transferred to 2 mL screw-cap tubes. 50 μL freshly prepared Lysozyme solution (20 mg/mL in TES, Amresco lyophilized powder, 23500 U/mg) was added to each resuspended pellet and incubated at room temperature for 15 minutes with continuous inversion. Sodium dodecyl sulfate (SDS) was added to a final concentration of 0.5% w/v and samples were incubated at room temperature for 10 minutes with continuous inversion. Samples were pelleted and the volume was reduced to 400 μL. 50 μL 10X Lysis Buffer (100 mM Tris pH 7.5, 100 mM NaCl, 1% IGEPAL CA-630 v/v) was added to each sample, followed by 50 μL freshly prepared 10X protease inhibitor (Roche cOmplete mini EDTA-free tablets). Cells were resuspended by pipetting and incubated on ice for 15 minutes. Manual lysis of cells was carried out by adding 400 μL 0.5 mm sterile glass beads to each tube and vortexing at maximum Hz for 30 seconds, followed by 30 seconds incubation on ice. Vortexing and ice incubation was repeated for 10 cycles. Bead-beaten samples were allowed to settle upright on ice for 15 minutes, then the liquid supernatant (~250 μL) was transferred to a new 1.5 mL tube. Sample volume was equilibrated to 500 μL with cold 2X NEBuffer 1.1 and incubated at 50° C for 10 minutes. After incubation, 30 μL 10% Triton X-100 v/v was added to each tube, mixed by inversion. Cross-linked DNA fragments were digested overnight with 50 U Sau3AI. Digested DNA complexes were pelleted (20,000 g, 4° C, 5 min.), gently washed with cold 1X NEBuffer 2, and resuspended in 200 μL NEBuffer 2.

Digested DNA was heated to 50° C for 5 minutes to melt paired sticky ends then put into a 200 μL Klenow fragment (exo-, NEB) fill-in reaction containing 36 μM biotin-14-dCTP (Thermo Fisher) and equimolar amounts of dATP, dTTP, and dGTP. Reactions were carried out for 2 hours at room temperature and the polymerase was quenched by adding EDTA to a final concentration of 10 mM. The full volume of each fill-in reaction was put into a dilute blunt-end ligation reaction (640 U T4 DNA Ligase, NEB) and allowed to incubate overnight at 15° C. Protein and crosslink digestion was carried out by adding 50 μL freshly prepared 20 mg/mL Proteinase K (VWR, freeze-dried powder suspended in 10 mM Tris, 1 mM MgCl_2_, 50% glycerol, pH 7.5) and incubating at 65° C for 6 hours. This digestion was repeated once. Protein was removed by phenol:chloroform extraction and ligated DNA was precipitated from the aqueous fraction with one volume 5M ammonium acetate and 4 volumes cold absolute ethanol. Clean DNA was quantified, and at least 1 μg but no more than 5 μg DNA was put into an end-resection reaction (5 U T4 DNA Polymerase, NEB) to remove biotin from unligated ends. Exonuclease activity of the polymerase was quenched with 5 mM EDTA and free biotinylated nucleotides were removed via 1.8X Ampure XP bead cleanup. Biotinylated DNA was immobilized on M280 streptavidin beads using the Invitrogen kilobaseBINDER Kit. Bead-bound DNA was quantified and prepared for sequencing using Illumina’s Nextera XT kit. Multiplexed libraries were size-selected with Ampure XP beads, quantified, and pooled for sequencing on an Illumina NextSeq 2×150 paired-end platform.

### Mock Community Methods

*Bacillus subtilis* containing pDR244, *Pseudomonas putida* containing pKJK5, and *Escherichia coli* containing RP4 were cultured in LB under antibiotic selection to maintain plasmids (spectinomycin, tetracycline, and kanamycin, respectively). Overnight cultures were washed with PBS, resuspended in PBS + 20% glycerol v/v, and frozen as aliquots, with one aliquot of each retained for titer determination on selective agar media. To create the mock community, 5×10^8^ colony forming units from each frozen stock was thawed and combined, and immediately carried through formaldehyde crosslinking as described for stool. Mock community Hi-C sequences were mapped with HiC-Pro against reference genomes and plasmids using default settings. Valid pairs, *i.e*. those that map to different restriction fragments, were compartmentalized into groups based on whether or not they connected the genome-genome, genome-plasmid, or plasmid-plasmid and coded according to the expected plasmid-host relationship.

### Quality Filtering and Assembly

Metagenomic and Hi-C sequences were quality filtered using Prinseq^50^ v0.20.2 to derepelicate, Bmtagger^51^ (v2/21/14) to remove human reads, and Trimmomatic^52^ v0.36 to remove adapters and quality filter reads (using settings: Leading:3, Trailing:3, Slidingwindow 4:15, Minlen: 50). Metagenomic reads were assembled using SPAdes^53^v3.13.2 with ‘-meta’ setting with a minimum contig size of 1,000bp. Genes on these contigs were called using Prodigal^54^ v2.6.3. PhageFinder^55^ v2.1 was used to identify large prophage regions and were excised from the first to the last phage gene called and considered separate contigs, unless the surrounding regions were less than 1,000bp, in which case they were also included as the excised phage.

### Metagenomic Binning

Contigs were binned using several tools (Maxbin^56^, MetaBat^57^, and Concoct^58^), culminating with a metagenomic binning aggregation strategy, DAS Tool^59^, we assessed genome contamination using CheckM and removed bins with contamination >10%, resulting in quality metagenomic bins although in many cases partial bins. To prevent overcalling of partial bins, downstream analyses aggregate at the taxon rather than individually calling unique bins.

### Taxonomic Identification

Kraken was applied to each metagenomic bin and annotated each contig individually using its algorithm. Then we assigned each bin the lowest taxonomic level at which more then 50% of the bin was assigned by Kraken with contigs weighted by length (bp). Contigs assigned Eukaryotic taxonomies were removed from further analysis.

### Antibiotic Resistance Genes

All contigs were annotated with CARD’s (Comprehensive Antibiotic Resistance Database) Resistance Gene Identifier (RGI)^60^ 3.2.1 against the CARD^61^ database and with HMMer^62^ against the Resfams^63^ database with the gathering cutoff. AR genes were clustered using CD-HIT-EST^64^ (identity:0.99; word size:8; length difference cutoff: 0.9) after they were sorted by length. Antibiotic resistance mechanisms are defined in Extended Data Table 4. We focused on AR genes that are commonly harbored by the most problematic MDR bacteria^65^ and that confer resistance to antibiotics that are most frequently relied on in neutropenic patients:

| Drug | Resistance determinant |
| --- | --- |
| Oxacillin<sup>66</sup> | *mecA*, *mecC* |
| Penicillin<sup>67,68,69</sup> | *pbp2b*, *pbp2x* |
| Ampicillin<sup>70</sup> | *bla*<sub>TEM</sub>, *bla*<sub>SHV</sub> |
| Cephalosporin<sup>71,72</sup> | *bla*<sub>TEM</sub>, *bla*<sub>SHV</sub>, *bla*<sub>CTX-M</sub>,<br>*bla*<sub>CMY</sub>, *bla*<sub>MIR</sub>, *bla*<sub>MOX</sub>, *bla*<sub>LAT</sub>, *bla*<sub>FOX</sub>, *bla*<sub>DHA</sub>, *bla*<sub>ACT</sub>, *bla*<sub>CFE</sub> |
| Carbapenem<sup>73</sup> | *bla*<sub>KPC</sub>, *bla*<sub>NDM</sub>, *bla*<sub>VIM</sub>, *bla*<sub>IMP</sub>, *bla*<sub>OXA-48</sub>, *bla*<sub>OXA-23</sub>, *oprD* |
| Fluoroquinolones<sup>74</sup> | *gyrA*, *gyrB*, *parC*, *parE*, *qnrA*, *qnrS* (Note: we considered any *qnr* gene.) |
| Aminoglycosides<sup>75</sup> | *aac*(3'), *aac*(6'), *aad* |

Genes in bold above represent those genes that were identified in our cohort’s microbiomes.

### Mobile Genetic Element annotation

All contigs and excised prophage contigs were assessed for the presence of mobile genes using several programs. Contigs were mapped using BLASTN to PlasmidFinder^76^ database (best hit, minimum 80% identity and 60% coverage), NCBI’s genomic plasmids downloaded (05/10/2017) (best hit, minimum 1000bp, minimum 80% identity), and IMMEdb^77^ (best hit, min 1,000 bp and 80% identity). Contigs were also identified as plasmids using PlasFlow^78^ with threshold of 0.95. Genes were mapped using BLASTP to ACLAME^79^ database v0.4 (besthit, min 80% identity and 60% coverage) and PHASTER^80^ prophage/virus database (v8/3/17) (best hit, min 80% identity and 60% coverage). Genes were mapped using HMMER^81^ v3.1b2 to Pfam^82^ and known plasmid, phage, and transposons were identified^83,84^. A search of common mobile gene terms against Pfam descriptions was carried out. Terms included for transposon: transpos, insertion element, is element, IS[0-9]; phage: phage, tail protein, tegument, capsid, relaxase, tail fibre, tail assembly, tail sheath, tail tube; plasmid: conjug, Trb, type IV, Tra[A-Z], mob, Vir[A-Z][0-9], t4ss, resolvase, plasmid; other: integrase. All Pfam IDs and descriptions are listed in the Extended Data Table 5. Contigs were also annotated for insertion sequence (IS) elements using ISEScan^85^ v1.5.4. Contigs with taxonomies assigned to the ‘Virus’ domain were considered phage. Contigs with any mobile annotation were annotated as MGEs.

### Sequence Mapping

Paired-end metagenomic sequences were mapped to the metagenomic contigs using BWA-MEM^86^ v0.7.13 requiring primary only alignments and filtered at 90% identity. Paired-end metagenomic sequences were also mapped separately to the AR and mobile gene clusters and filtered at 99% identity. Contig and individual AR and mobile gene RPKM values were calculated using mapped metagenomic reads (total reads mapped to the contigs with >80% contig coverage, divided by the length of the contigs per kilobase and the total read count in that sample per million). Hi-C reads were mapped with HiC-Pro using default parameterswhich internally uses Bowtie2. HicPro requires valid pairs to map to different restriction fragments and allows only unique mapping of reads.

### Cleanliness comparisons

Reads mapping between two different contigs were included in the analysis if neigher contig carried a mobile gene. Taxonomic associations were determined from residency in a metagenomic bin annotated to at least the taxonomic level of interest.

### Mobile and AR gene associations

All mobile and AR gene-containing contigs, including excised phage, were associated with taxa if they were linked to a Hi-C clustered genomic contig with at least two Hi-C read pairs or if they were clustered into an annotated genomic bin. Hi-C linkages between MGEs and their genomic bins are more robust if Hi-C reads map more closely (*i.e.* smaller linear distance (bp)) with the genes that are annotated as mobile. To assess this, we calculated the genetic distance between the mobile or AR gene and the nearest Hi-C read linking any contig with a particular taxonomy. Often this resulted in multiple linkages between the mobile contig and taxonomic contigs clustered with the same taxa. We therefore assessed the strongest data linking the two, the minimum genetic distance, considering the other reads as further support for this gene-taxon assignment.

### Comparison of HGT between individual taxa

First, we compared HGT observed between species (as shown in Figure 2B,C and Extended Data Figure 8), defined above, through Hi-C read pair linkages. To create an HGT network, we examined the number of unique (defined as 99% sequence identity) AR or mobile gene linked to genomic bins for each particular taxa. Consequently we could identify taxa-taxa connections based on these identified gene sharing events.

We assessed the rate of HGT per 100 species-species comparisons at different taxonomic levels within and between patients, as a comparison with Smillie et al. (2011)^87^. For comparisons between species, we compared each species within a single genus to one another. For every other taxonomic level, we compared species that differed according to that taxonomic level (*i.e*. for comparisons between families, species of one family, *e.g.* the Enterobacteriaceae, were compared exclusively with species in other taxonomic families). When comparing two species, we considered HGT events as those taxa sharing at least one gene of interest (AR or mobile gene) at >99% identity. We compared HGT within each patient or performed pairwise comparisons between the 9 individuals. For each taxonomic level, we compared within vs. between patients using a Mann-Whitney U-test.

### AR gene and phage machinery gene host specificity

Genes of interest associated with taxa through Hi-C alone (*i.e*. not including taxa originally assigned to a contig that contained that AR gene or phage machinery gene) were compared to taxonomies identified by comparison using BLASTN (e-value < 1e-100) to PATRIC^88^ genome database (downloaded October 1, 2018) for the AR genes or compared to NCBI nt database (downloaded June 4, 2018) for the phage machinery genes and placed into one of several categories defined in Figure 1. This was assessed at different taxonomic levels.

### Network density

Network density was calculated by dividing the number of observed connections between mobile or AR genes and binned organisms in each sample out of the theoretical maximum number of connections (number of AR genes or mobile genes multiplied by number of distinct organisms). Number of total species in a population were identified from MetaPhlan.

### Measuring novel HGT during individuals’ timecourses

Within an individuals’ timecourse, we identified novel HGT events by comparing the first timepoint to subsequent timepoints and requiring that the gene-taxa connection met several criteria. HGT events were only considered between donor organisms strongly linked with a mobile or mobile AR gene at the start of the timecourse and recipient organisms present, albeit unlinked to the mobile or mobile AR gene, at the start of the timecourse. We sequenced the initial timepoint 3-4 times more deeply than the remainder of the timecourse to be able to distinguish between migration of strains and HGT. Mobile or AR gene-containing contigs were required to be linked via at least 2 Hi-C reads (mean = 28.6) to genome assemblies that were taxonomically annotated at the level of genus or species. We required an absence of association between the mobile or AR gene of interest and potential recipient taxa. In other words, one Hi-C read was sufficient to disqualify a putative HGT event, as was any taxonomic marker on that contig associating it with a congruent recipient taxon, or any association with a genome assembly with any congruent higher-order taxonomy. We required recipient taxa to have at least 2 Hi-C reads (mean = 14.8) associating each mobile or AR gene-containing contig with the recipient genome assembly. We tallied the number of HGT events that were supported by more than one timepoint, both strictly by the same genes and also by the same taxa; as well as those that were supported across multiple contigs.

### Data Availability

Metagenomic and Hi-C sequences, filtered for quality and human-reads will be uploaded to NCBI’s Short Read Archive. Code relies heavily on published packages listed above. Custom code available upon request.

## Acknowledgements

This study was funded by the Centers for Disease Control (OADS BAA 2016-N-17812) and by the National Sciences Foundation (Awards #1661338 and #1650122). M.J.S. is funded by the NIAID (K23 AI114994). I.L.B. is funded by the NIH (1DP2HL141007-01) and is a Sloan Foundation Research Fellow, a Packard Fellowship in Science and Engineering, and a Pew Foundation Biomedical Scholar. Albert Vill is a Cornell University Centers for Vertebrate Genomics 2019 Distinguished Scholar.

## References

1. Li, J. et al. Antibiotic Treatment Drives the Diversification of the Human Gut Resistome. Genomics Proteomics Bioinformatics 17, 39–51 (2019).

2. Kaminski, J. et al. High-Specificity Targeted Functional Profiling in Microbial Communities with ShortBRED. PLoS Comput. Biol. 11, e1004557 (2015).

3. Sommer, M. O. A., Dantas, G., Church, G. M. Functional characterization of the antibiotic resistance reservoir in the human microflora. Science. 325, 1128–1131 (2009).

4. Satlin, M. J. & Walsh, T. J. Multidrug-resistant Enterobacteriaceae, Pseudomonas aeruginosa, and vancomycin-resistant Enterococcus: Three major threats to hematopoietic stem cell transplant recipients. Transpl Infect Dis 19,(2017).

5. Huddleston, J. R. Horizontal gene transfer in the human gastrointestinal tract: potential spread of antibiotic resistance genes. Infect Drug Resist. 7, 167–176. (2014).

6. Satlin, M. J. & Walsh, T. J. Multidrug-resistant Enterobacteriaceae, Pseudomonas aeruginosa, and vancomycin-resistant Enterococcus: Three major threats to hematopoietic stem cell transplant recipients. Transpl Infect Dis 19,(2017).

7. Sommer, M. O. A., Dantas, G., Church, G. M. Functional characterization of the antibiotic resistance reservoir in the human microflora. Science. 325, 1128–1131 (2009).

8. Gibson, M. K., Forsberg, K. J. & Dantas, G. Improved annotation of antibiotic resistance determinants reveals microbial resistomes cluster by ecology. ISME J 9, 207–216 (2015).

9. Kaminski, J. et al. High-Specificity Targeted Functional Profiling in Microbial Communities with ShortBRED. PLoS Comput. Biol. 11,e1004557 (2015).

10. Yaffe, E. & Relman, D. A. Tracking microbial evolution in the human gut using Hi-C reveals extensive horizontal gene transfer, persistence and adaptation. Nat Microbiol (2019) doi:10.1038/s41564-019-0625-0.

11. Marbouty, M. et al. Metagenomic chromosome conformation capture (meta3C) unveils the diversity of chromosome organization in microorganisms. Elife 3,e03318 (2014).

12. Burton, J. N., Liachko, I., Dunham, M. J. & Shendure, J. Species-level deconvolution of metagenome assemblies with Hi-C-based contact probability maps. G3 (Bethesda) 4, 1339–1346 (2014).

13. Beitel, C. W. et al. Strain- and plasmid-level deconvolution of a synthetic metagenome by sequencing proximity ligation products. PeerJ 2,e415 (2014).

14. Marbouty, M. et al. Metagenomic chromosome conformation capture (meta3C) unveils the diversity of chromosome organization in microorganisms. Elife 3,e03318 (2014).

15. Stewart, R. D., et al. Assembly of 913 microbial genomes from metagenomic sequencing of the cow rumen. Nature Comm 9, 870 (2018).

16. Bickhart, D. M. et al. Assignment of virus and antimicrobial resistance genes to microbial hosts in a complex microbial community by combined long-read assembly and proximity ligation. Genome Biology 20, 153 (2019).

17. Stalder, T., Press, M. O., Sullivan, S., Liachko, I. & Top, E. M. Linking the resistome and plasmidome to the microbiome. The ISME Journal 13, 2437–2446 (2019).

18. Marbouty, M., Baudry, L., Cournac, A. & Koszul, R. Scaffolding bacterial genomes and probing host-virus interactions in gut microbiome by proximity ligation (chromosome capture) assay. Sci Adv 3,e1602105 (2017).

19. Satlin, M. J. & Walsh, T. J. Multidrug-resistant Enterobacteriaceae, Pseudomonas aeruginosa, and vancomycin-resistant Enterococcus: Three major threats to hematopoietic stem cell transplant recipients. Transpl Infect Dis 19,(2017)

20. Pop, M. Genome assembly reborn: recent computational challenges. Brief Bioinform. 10, 354–366 (2009).

21. Krawczyk, P. S., Lipinski, L., Dziembowski, A. PlasFlow: predicting plasmid sequences in metagenomic data using genome signatures. Nucleic Acids Res. 46, e35 (2018). Apr 6; 46(6): e35.

22. Wu, Y.-W., Simmons, B. A. & Singer, S. W. MaxBin 2.0: an automated binning algorithm to recover genomes from multiple metagenomic datasets. Bioinformatics 32, 605–607 (2016).

23. Sieber, C. M. K. et al. Recovery of genomes from metagenomes via a dereplication, aggregation and scoring strategy. Nat Microbiol. 3, 836–843. (2018)

24. Pop, M. Genome assembly reborn: recent computational challenges. Brief Bioinform. 10, 354–366 (2009).

25. Ross, A., Ward, S. & Hyman, P. More Is Better: Selecting for Broad Host Range Bacteriophages. Front. Microbiol. 7,(2016).

26. Yu, J., Lim, J.-A., Kwak, S.-J., Park, J.-H. & Chang, H.-J. Comparative genomic analysis of novel bacteriophages infecting Vibrio parahaemolyticus isolated from western and southern coastal areas of Korea. Arch. Virol. 163, 1337–1343 (2018).

27. Doulatov, S. et al. Tropism switching in Bordetella bacteriophage defines a family of diversity-generating retroelements. Nature 431, 476–481 (2004).

28. Ross, A., Ward, S. & Hyman, P. More Is Better: Selecting for Broad Host Range Bacteriophages. Front Microbiol 7, 1352 (2016).

29. Crémazy, F. G. et al. Determination of the 3D genome organization of bacteria using Hi-C. Methods Mol Biol. 1837, 3–18 (2018).

30. Marbouty, M. et al. Metagenomic chromosome conformation capture (meta3C) unveils the diversity of chromosome organization in microorganisms. Elife 3, e03318 (2014).

31. Brito, I. L. et al. Mobile genes in the human microbiome are structured from global to individual scales. Nature 535, 435–439 (2016).

32. Brito, I. L. et al. Mobile genes in the human microbiome are structured from global to individual scales. Nature 535, 435–439 (2016).

33. Yassour, M. et al. Natural history of the infant gut microbiome and impact of antibiotic treatments on strain-level diversity and stability. Sci Transl Med 8, 343ra81 (2016).

34. Smillie, C. S. et al. Ecology drives a global network of gene exchange connecting the human microbiome. Nature 480, 241–244 (2011).

35. Kaakoush, N. O. Insights into the Role of Erysipelotrichaceae in the Human Host. Front Cell Infect Microbiol. 5, 84 (2015).

36. Rossi, O. et al. Faecalibacterium prausnitzii A2-165 has a high capacity to induce IL-10 in human and murine dendritic cells and modulates T cell responses. Scientific Reports 6, 18507 (2016).

37. Zhu, C. et al. Roseburia intestinalis inhibits interleukin-17 excretion and promotes regulatory T cells differentiation in colitis. Mol Med Rep. 17, 7567–7574 (2018).

38. Modi, S. R., Lee, H. H., Spina, C. S. & Collins, J. J. Antibiotic treatment expands the resistance reservoir and ecological network of the phage metagenome. Nature 499, 219–222 (2013).

39. Diard, M. et al. Inflammation boosts bacteriophage transfer between Salmonella spp. Science 355, 1211–1215 (2017).

40. Bakkeren, E. et al. Salmonella persisters promote the spread of antibiotic resistance plasmids in the gut. Nature 573, 276–280 (2019).

41. Sommer, M. O. A., Dantas, G. & Church, G. M. Functional characterization of the antibiotic resistance reservoir in the human microflora. Science 325, 1128–1131 (2009).

42. Bakkeren, E. et al. Salmonella persisters promote the spread of antibiotic resistance plasmids in the gut. Nature 573, 276–280 (2019).

43. Bertrand, D. et al. Hybrid metagenomic assembly enables high-resolution analysis of resistance determinants and mobile elements in human microbiomes. Nat. Biotechnol. 37, 937–944 (2019).

44. Knöppel, A., Lind, P. A., Lustig, U., Näsvall, J. & Andersson, D. I. Minor fitness costs in an experimental model of horizontal gene transfer in bacteria. Mol. Biol. Evol. 31, 1220–1227 (2014).

45. McCarthy, A. J. et al. Extensive horizontal gene transfer during Staphylococcus aureus co-colonization in vivo. Genome Biol Evol 6, 2697–2708 (2014).

46. Citorik, R. J., Mimee, M., & Lu, T. K. Sequence-specific antimicrobials using efficiently delivered RNA-guided nucleases. Nature Biotechnology. 32, 1141–1145 (2014).

47. Vercoe, R. B. et al. Cytotoxic chromosomal targeting by CRISPR/Cas systems can reshape bacterial genomes and expel or remodel pathogenicity islands. PLoS Genet. 9, e1003454 (2013).

48. Yosef, I., Manor, M., Kiro, R., & Qimron U. Temperate and lytic bacteriophages programmed to sensitize and kill antibiotic-resistant bacteria. Proc Natl Acad Sci U S A. 112, 7267–72 (2015).

49. Hi-C deconvolution of a human gut microbiome yields high-quality draft genomes and reveals plasmid-genome interactions. bioRxiv. doi:10.1101/1987713v1. (2017)

## Extended Data References

50. Schmieder, R. & Edwards, R. Quality control and preprocessing of metagenomic datasets. Bioinformatics 27, 863–864 (2011).

51. Rotmistrovsky, K. & Agarwala, R. BMTagger: Best Match Tagger for removing human reads from metagenomics datasets. Unpublished (2011).

52. Bolger, A. M., Lohse, M. & Usadel, B. Trimmomatic: a flexible trimmer for Illumina sequence data. Bioinformatics 30, 2114–2120 (2014).

53. Nurk, S., Meleshko, D., Korobeynikov, A. & Pevzner, P. A. metaSPAdes: a new versatile metagenomic assembler. Genome Res. 27, 824–834 (2017).

54. Hyatt, D. et al. Prodigal: prokaryotic gene recognition and translation initiation site identification. BMC Bioinformatics 11, 119 (2010).

55. Fouts, D. E. Phage_Finder: Automated identification and classification of prophage regions in complete bacterial genome sequences. Nucleic Acids Res 34, 5839–5851 (2006).

56. Wu, Y.-W., Simmons, B. A. & Singer, S. W. MaxBin 2.0: an automated binning algorithm to recover genomes from multiple metagenomic datasets. Bioinformatics 32, 605–607 (2016).

57. Kang, D. D. et al. MetaBAT 2: an adaptive binning algorithm for robust and efficient genome reconstruction from metagenome assemblies. PeerJ 7, e7359 (2019).

58. Alneberg, J. et al. Binning metagenomic contigs by coverage and composition. Nat Methods 11, 1144–1146 (2014).

59. Sieber, C. M. K. et al. Recovery of genomes from metagenomes via a dereplication, aggregation and scoring strategy. Nat Microbiol. 3, 836–843. (2018)

60. Jia, B. et al. CARD 2017: expansion and model-centric curation of the comprehensive antibiotic resistance database. Nucleic Acids Res 45,D566–D573 (2017).

61. McArthur, A. G. et al. The comprehensive antibiotic resistance database. Antimicrob. Agents Chemother. 57, 3348–3357 (2013).

62. Mistry, J., Finn, R. D., Eddy, S. R., Bateman, A. & Punta, M. Challenges in homology search: HMMER3 and convergent evolution of coiled-coil regions. Nucleic Acids Res. 41,e121 (2013).

63. Gibson, M. K., Forsberg, K. J. & Dantas, G. Improved annotation of antibiotic resistance determinants reveals microbial resistomes cluster by ecology. ISME J 9, 207–216 (2015).

64. Li, W. & Godzik, A. Cd-hit: a fast program for clustering and comparing large sets of protein or nucleotide sequences. Bioinformatics 22, 1658–1659 (2006).

65. Centers for Disease Control. Antibiotic Resistance Threats in the United States. Atlanta, GA (2013)

66. Lakhundi, S. & Zhang, K. Methicillin-Resistant Staphylococcus aureus: Molecular Characterization, Evolution, and Epidemiology. Clin Microbiol Rev. 31, e00020–18 (2018).

67. Magill, S. S. et al. Prevalence of antimicrobial use in US acute care hospitals, May-September 2011. JAMA 312, 1438–1446 (2014).

68. Dowson, C. G. et al. Penicillin-resistant viridans streptococci have obtained altered penicillin-binding protein genes from penicillin-resistant strains of Streptococcus pneumoniae. Proc Natl Acad Sci U S A. 87, 5858–5862 (1990).

69. van der Linden, M. et al. Insight into the Diversity of Penicillin-Binding Protein 2x Alleles and Mutations in Viridans Streptococci. Antimicrob Agents Chemother. 61,e02646–16 (2017).

70. Paterson, D. L. & Bonomo, R. A. Extended-Spectrum β-Lactamases: a Clinical Update. . Clinical Microbiology Reviews. 18, 657–686. (2005).

71. Paterson, D. L. & Bonomo, R. A. Extended-spectrum beta-lactamases: a clinical update. Clin Microbiol Rev. 18, 657–686. (2005).

72. Strahilevitz, J., Jacoby, G. A., Hooper, D. C., Robicsek, A. Plasmid-mediated quinolone resistance: a multifaceted threat. Clin Microbiol Rev. 22, 664–689 (2009).

73. Queenan, A. M., & Bush, K. Carbapenemases: the Versatile β-Lactamases. Clinical Microbiology Reviews. 20, 440–458 (2007).

74. Hooper, D. C., & Jacoby, G. A.Topoisomerase Inhibitors: Fluoroquinolone Mechanisms of Action and Resistance.Cold Spring Harb Perspect Med. 6, a025320 (2016).

75. Doi, Y., Wachino, J. I., Arakawa, Y. Aminoglycoside Resistance: The Emergence of Acquired 16S Ribosomal RNA Methyltransferases. Infect Dis Clin North Am. 30, 523–537 (2016).

76. Carattoli, A. et al. In silico detection and typing of plasmids using PlasmidFinder and plasmid multilocus sequence typing. Antimicrob. Agents Chemother. 58, 3895–3903 (2014).

77. Jiang, X., Hall, A. B., Xavier, R. J., & Alm, E. J. Comprehensive analysis of mobile genetic elements in the gut microbiome reveals phylum-level niche-adaptive gene pools BioRxiv. doi:10.1101/214213, (2017)

78. Krawczyk, P. S., Lipinski, L. & Dziembowski, A. PlasFlow: predicting plasmid sequences in metagenomic data using genome signatures. Nucleic Acids Res. 46, e35 (2018).

79. Leplae, R., Lima-Mendez, G. & Toussaint, A. ACLAME: a CLAssification of Mobile genetic Elements, update 2010. Nucleic Acids Res. 38,D57–61 (2010).

80. Arndt, D. et al. PHASTER: a better, faster version of the PHAST phage search tool. Nucleic Acids Res. 44, W16–21 (2016).

81. Mistry, J., Finn, R. D., Eddy, S. R., Bateman, A. & Punta, M. Challenges in homology search: HMMER3 and convergent evolution of coiled-coil regions. Nucleic Acids Res. 41, e121 (2013).

82. Finn, R. D. et al. The Pfam protein families database: towards a more sustainable future. Nucleic Acids Res. 44, D279–285 (2016).

83. Sczyrba, A. et al. Critical Assessment of Metagenome Interpretation—a benchmark of metagenomics software. Nature Methods 14, 1063–1071 (2017).

84. Schlüter, A., Krause, L., Szczepanowski, R., Goesmann, A. & Pühler, A. Genetic diversity and composition of a plasmid metagenome from a wastewater treatment plant. Journal of Biotechnology 136, 65–76 (2008).

85. Xie, Z. & Tang, H. ISEScan: automated identification of insertion sequence elements in prokaryotic genomes. Bioinformatics 33, 3340–3347 (2017).

86. Li, H. & Durbin, R. Fast and accurate short read alignment with Burrows-Wheeler Transform. Bioinformatics, 25: 1754–60. (2009).

87. Smillie, C. S. et al. Ecology drives a global network of gene exchange connecting the human microbiome. Nature 480, 241–244 (2011).

88. Wattam, A. R. et al. PATRIC, the bacterial bioinformatics database and analysis resource. Nucleic Acids Res. 42, D581–591 (2014).

